# Phenotype-Centric Modeling for Rational Metabolic Engineering

**DOI:** 10.1101/2021.11.26.470163

**Authors:** Miguel Á. Valderrama-Gómez, Michael A. Savageau

## Abstract

Phenotype-centric modeling enables a paradigm shift in the analysis of kinetic models. It brings the focus to a network’s biochemical phenotypes and their relationship with measurable traits (e.g., product yields, system dynamics, signal amplification factors, etc.) and away from computationally intensive parameter sampling and numerical simulation. Here, we explore applications of this new modeling strategy in the field of Rational Metabolic Engineering using the amorphadiene biosynthetic network as a case study. Our phenotype-centric approach not only identifies known beneficial intervention strategies for this network, but it also provides an understanding of mechanistic context for the validity of these predictions. Additionally, we propose a set of hypothetical strains with the potential to outperform reported production strains and enhance the mechanistic understanding of the amorphadiene biosynthetic network. We believe that phenotype-centric modeling can advance the field of Rational Metabolic Engineering by enabling the development of next generation kinetics-based algorithms and methods that do not rely on *a priori* knowledge of kinetic parameters but allow a structured, global analysis of the design space of parameter values.

## 1. Introduction

Metabolic Engineering aims at developing cellular factories to produce valuable chemicals by altering the metabolism of microbial strains through genetic engineering (Bailey, 1991; Lee et al., 2012; Chubukov et al., 2016). During the last decades, computational methods have enabled the discovery of non-intuitive strategies enhancing the production of a variety of target molecules (Nakamura and Whited, 2003; Lee et al., 2005; Yim et al., 2011; Paddon et al., 2013; Harder et al., 2016), giving rise to a model-based Metabolic Engineering that is increasingly less dependent on experimental intuition. Existing mathematical frameworks for rational Metabolic Engineering typically fall within one of two categories: kinetics- (mechanistic) or constraint-based methods (Valderrama-Gómez et al., 2017). The former is the gold standard and has the potential to capture intricate interactions between different levels of cellular organization (transcription, translation, and metabolism), which need to be rigorously integrated to understand and successfully optimize biological systems. Mechanistic models are truly predictive because they provide a rigorous link between metabolite concentrations, enzyme availability, and intracellular flux distributions. However, the application of mechanistic modeling in Metabolic Engineering has been rather limited, mainly due to a bottleneck caused by the large number of associated parameters with unknown values (Chowdhury, 2015), and structural uncertainties arising from unknown molecular interactions (Link et al., 2014). Consequently, constraint-based modeling (e.g., flux balance analysis) has been the method of choice to rationally guide the development of production strains (Zomorrodi, 2012; Valderrama-Gómez et al., 2017), which is reflected in most strain design algorithms being based on stoichiometric descriptions of cellular metabolism (Valderrama-Gómez et al., 2017).

Even though constraint-based modeling has proven to be useful in characterizing certain aspects of metabolic networks (e.g., maximal theoretical yields), important features such as network dynamics and metabolite concentrations are outside the scope of these models (Wiechert and Noack, 2011). Thus, implementing engineering strategies suggested by constraint-based metabolic models can potentially lead to non-viable strains because critical aspects of the metabolic system, such as dynamical instability and concentrations of potentially toxic metabolite intermediates are not considered.

Overcoming the limitations of kinetic modeling will require a radical change in how these models are formulated and analyzed. In the last decade, ensemble modeling of metabolic networks has emerged as a useful approach to address both parametric (Tran et al., 2008; Lee et al., 2014) and structural (Link et al., 2014) uncertainties in kinetic models. Instead of analyzing a single model, this approach considers thousands of models, each exhibiting a different set of parameter values or alternative molecular interactions. A recently developed *phenotype-centric* modeling strategy (Lomnitz and Savageau, 2016; Valderrama-Gómez et al., 2018) offers enormous potential for the field of rational Metabolic Engineering by allowing the analysis of mechanistic models without *a priori* knowledge of kinetic parameters (Valderrama-Gómez, et al., 2020). The strategy combines a model decomposition technique with linear programming in logarithmic space to identify a space-filling set of *biochemical phenotypes*, each one valid within a high-dimensional polytope in the design space of parameter values. Biochemical phenotypes are mathematically described by a simplified set of S-system differential equations (i.e., sub-systems), and an accompanying set of linear inequalities in logarithmic space. The mathematical object defining a biochemical phenotype involves all the system variables, parameters, and non-dominant processes. This is critically important because the ‘dominance concept’ may erroneously suggest a model reduction method with significant losses. It follows that any point in the parameter space is contained within at least one such biochemical phenotype. Powerful mathematical techniques have been developed to characterize S-systems in terms of steady states, signal amplification factors (logarithmic gains), phenotypic volumes, and dynamic behavior (Savageau et al., 2009; Fasani and Savageau, 2010; Lomnitz and Savageau, 2016). These features link experimentally observable biological phenotypes with specific regions in the parameter space to produce a finite, chunked and highly structured System Design Space.

In a recent work, we briefly discussed the potential of the phenotype-centric approach in Metabolic Engineering by analyzing the protocatechuate metabolic system of *Acinetobacter* (Valderrama-Gómez et al., 2020). We built a mathematical model considering the transport of protocatechuate into the cell and its subsequent enzymatic degradation, the synthesis of the intervening enzymes and, a signaling layer controlling the synthesis of mRNA molecules. The mathematical model encompassed 30 parameters whose values were assumed to be unknown. Using the phenotype-centric approach, we were able to identify a biochemical phenotype and values for all system parameters that potentially correspond to the natural operating point of the system. Moreover, we proposed several engineering strategies to increase the pathway flux without increasing the intracellular concentration of toxic pathway intermediates.

In this work, we will further explore the utility of the phenotype-centric strategy in the field of Metabolic Engineering. We will use the amorphadiene biosynthetic network shown in Fig. 1 as a case study. Weaver et. al. (2015) mechanistically characterized this metabolic system in *Escherichia coli* using a kinetic model to identify various engineering strategies to increase productivity. *In silico* predictions were experimentally implemented and the performance of the resulting engineered strains closely matched model predictions. The work by Weaver et. al. (2015) will help us contrast traditional strain optimization procedures using kinetic models in a simulation-centric approach with our novel phenotype-centric strategy. As we will show later, our analysis not only reproduces the predictions made by Weaver et. al. (2015) but also provides a structured context in the System Design Space for which those predictions are valid. Furthermore, engineering strategies covering different regions in Design Space are also identified by our approach in a computationally efficient way that does not involve parameter sampling or numerical solution of the underlying kinetic model.

**Figure 1.**
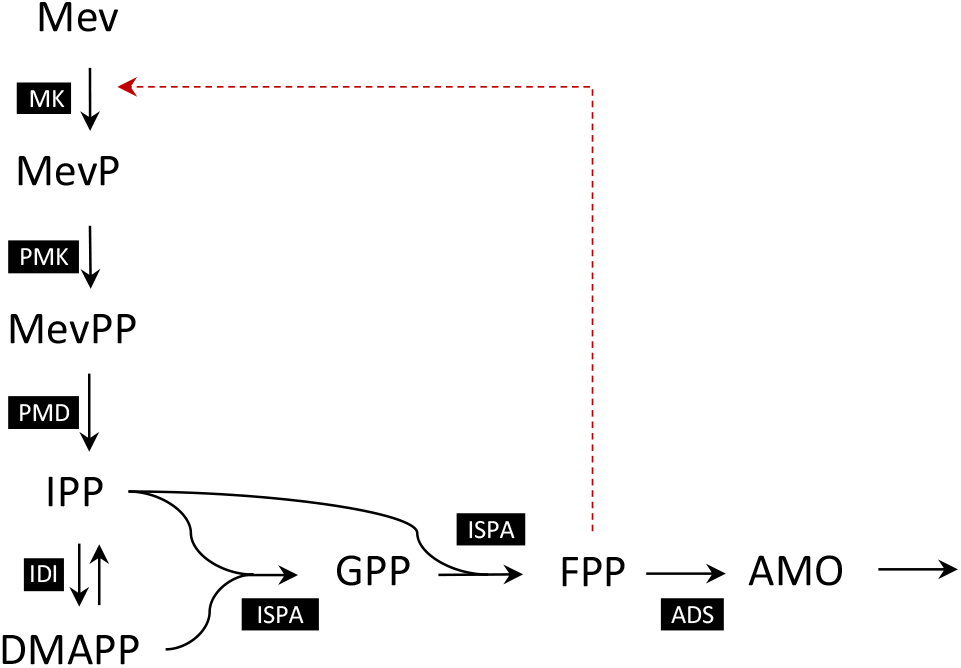
The Amorphadiene Biosynthetic Pathway. Amorphadiene is synthesized from externally supplied mevalonate in a series of seven enzyme-catalyzed reactions that involve the interaction of eight different metabolites. Enzymes are represented by black boxes using the following abbreviations: *MK*: Mevalonate kinase, *PMK*: Phosphomevalonate kinase, *PMD*: Mevalonate diphosphate decarboxylase, *IDI*: Isopentenyl-diphosphate isomerase, *ISPA*: Farnesyl pyrophosphate synthase, *ADS*: Amorphadiene synthase. Metabolites are represented by the following abbreviations: *Mev*: Mevalonate, *MevP*: Mevalonate phosphate, *MevPP*: Mevalonate pyrophosphate, *IPP*: Isopentenyl pyrophosphate, *DMAPP*: Dimethylallyl pyrophosphate, *GPP*: geranyl diphosphate, *FPP*: farnesyl diphosphate, *AMO*: Amorphadiene. The isomerization reaction of IPP and DMAPP catalyzed by IDI is the only reversible reaction in the network. The pathway involves feedback inhibition of MK, the first enzyme, by the last metabolic intermediate, FPP.

We start in Section 2 by providing a description of the amorphadiene biosynthetic pathway and briefly summarizing previous findings by Weaver et. al. (2015). Relevant features of the phenotype-centric approach will be presented in Section 3. For a more detailed review of the method’s mathematical background along with its computational implementation, the interested reader is directed to previous publications (Savageau et al., 2009; Fasani and Savageau, 2010; Lomnitz and Savageau, 2016; Valderrama-Gómez et al., 2018; Valderrama-Gómez et al., 2020). Section 4 will illustrate different applications of the phenotype-centric strategy in Metabolic Engineering. Lastly, we conclude with a discussion and provide future directions in Section 5.

## 2. The Amorphadiene Biosynthetic Pathway

Amorphadiene is a volatile terpene involved in the synthesis of the anti-malarial drug artemisinin and its heterologous production was first reported in *E. coli* (Newman et al., 2006). Weaver et. al. (2015) mechanistically characterized a metabolic system which used externally provided mevalonate to synthesize amorphadiene in a series of seven enzyme-catalyzed steps. The authors employed a kinetic model to identify the concentration of amorphadiene synthase (ADS), the last enzyme of the pathway (Fig. 1), as one of the main engineering targets to improve productivity. The analysis of the kinetic model also showed that alleviating the feedback inhibition of mevalonate kinase (the first enzyme in the pathway) by farnesyl pyrophosphate (the final pathway intermediate) did not increase amorphadiene productivity, as previously hypothesized (Weaver et. al., 2015). Both predictions were based on a sensitivity analysis of the model parameters and were experimentally verified by constructing and characterizing three different strains: mbis3 (the base strain), saMK, and 10kADS. The kinetic model developed by Weaver et al. (2015) was based on a considerable body of previous experimental work that involved extracting values for 26 kinetic parameters from the literature and experimentally determining protein concentrations for all three strains. We slightly modified this model to consider a first-order output flux for amorphadiene (Eqs. S1 to S8). This modification does not affect the dynamics of the network’s metabolic pools and solely serves to conveniently characterize the flux through the pathway in the context of the phenotype-centric approach.

## 3. Materials and Methods

The System Design Space (Savageau et al., 2009) represents the mathematical foundation upon which the phenotype-centric modeling strategy (Valderrama-Gómez et al. 2018) is built. Here, we will provide a brief overview of key concepts as well as instructions to access computational tools to reproduce the results presented in this work.

### 3.1 System Design Space: Key Concepts

Biochemical systems described by the power-law functions of chemical kinetics and the rational functions of biochemical kinetics can be represented in a Generalized Mass Action (GMA) form (Savageau and Voit, 1987):

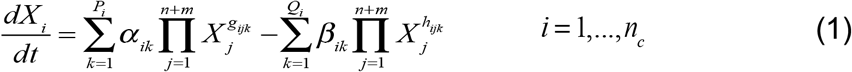

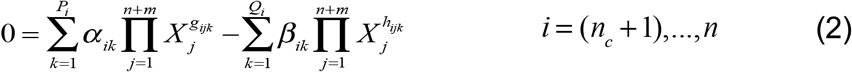

Where *α*_*ik*_ and *β*_*ik*_ represent rate constants, while *g*_*ijk*_ and *h*_*ijk*_ are kinetic orders. *P*_*i*_ and *Q*_*i*_ are the number of positive and negative terms in the *i*-th equation, respectively. Additionally, *X*_*i*_ represents the concentration of a chemical species in a system containing a total of *n* dependent and *m* independent variables. The set *n*_*c*_ of chemical variables represents all the chemical/biological entities (e.g., enzymes, metabolites, mRNA molecules, etc.) of the system. On the other hand, the set *n* − *n*_*c*_ contains auxiliary variables generated when recasting the system of ordinary differential equations into its GMA form. Environmental input variables for which a differential equation or algebraic constraint are not defined are treated as parameters. For any system in steady state, one of the positive and one of the negative terms will dominate over the others in each one of the *n* equations in the system. This gives rise to a so-called *dominant S-System* (Savageau, 1969; Savageau et al., 2009), which can be generically described by Eqs. 3 and 4:

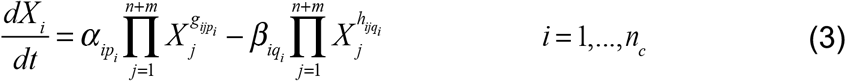

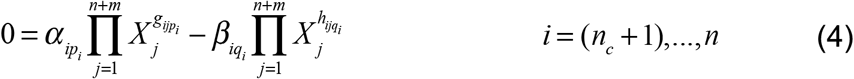

with *p*_*i*_ and *q*_*i*_ representing the indices of the dominant positive and dominant negative terms in the *i*-th equation, respectively. The validity of the dominant S-System implies certain conditions (Savageau et al., 2009; Fasani and Savageau, 2010), which are represented by inequalities of the form

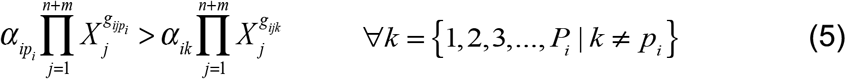

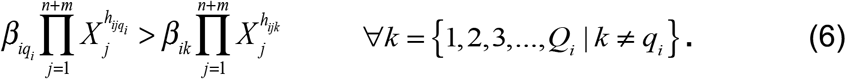

Here, *k* represents indices of corresponding non-dominant terms. Steady state concentrations of the dependent variables can be obtained by rearranging Eqs. 3 and 4 and taking logarithms on both sides:

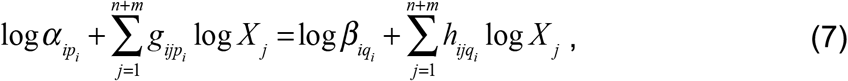

which can be written in matrix form as:

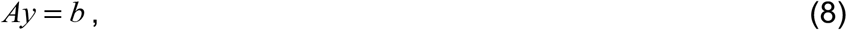

Where 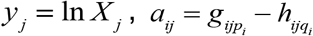 and *b*_*i*_ = ln(*β*_*in*_ / *α*_*iq*_). In a following step, dependent (*y*_*D*_) and independent (*y*_*I*_) variables are split to obtain:

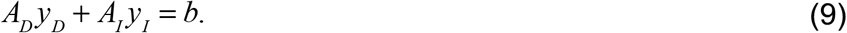

The vector of dependent concentration variables *y*_*D*_ can be obtained by matrix operations:

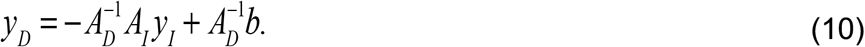

The flux through each metabolic pool can be obtained by a secondary matrix operation (Savageau, 2009). Because some have confused S-system equations in two different contexts, it should be noted that the original S-system equations were found in the context of a local (Taylor series) representation in logarithmic space and involved *real-valued* exponents (Savageau, 1969; 2009), whereas the S-system equations found in the global context of the System Design Space involve positive *integer-valued* exponents defined by the underlying chemical and biochemical kinetic mechanisms (Savageau et al. 2009). In any case, they have the same mathematical form, which makes them amenable to the same set of powerful linear methods.

#### 3.1.1 Biochemical Phenotypes

The concept of *biochemical phenotype* (or simply *phenotype*) is an integral element of the Design Space formalism and will be broadly used throughout this work. A phenotype is defined in the context of a mechanistic model of a biological system. The mathematical representation of a biochemical phenotype is given by a set of dominant S-system equations (Eqs. 3 and 4) and associated boundaries, which involve a comprehensive integration of information for all the system’s concentrations, fluxes, and parameters (Savageau et al., 2009; Fasani and Savageau, 2010). From a biological point of view, most of the mathematical properties of a biochemical phenotype, such as its logarithmic gains and dynamic behavior, can be experimentally measured, thus rendering biochemical phenotypes a powerful tool to design and optimize biochemical systems. Biochemical phenotypes can be categorized into two groups: pathological and physiological. The former is characterized by internal metabolic imbalances that result in the continual accumulation or depletion of at least one metabolic pool (Valderrama-Gómez et al. 2020). Conversely, physiological phenotypes exhibit at least one steady state, which can be either stable or unstable. Phenotypes have an associated case number and a signature that implies a specific set of dominance conditions.

#### 3.1.2 Logarithmic Gains

Logarithmic gains are amplification factors relating changes in input signals (independent variables, *y*_*I*_) to the resulting changes in output signals (dependent variables, *y*_*D*_) and are denoted by the symbol *L(y*_*D*_, *y*_*I*_). Strictly speaking, the term parameter sensitivity is used instead of logarithmic gain when the effect of varying a parameter on a dependent variable is analyzed. Both logarithmic gains and parameter sensitivities are properties that depend exclusively on the kinetic orders of the system and can be calculated for concentrations or fluxes (Savageau, 1971). They are valid throughout the entire polytope of a given biochemical phenotype. For simplicity, we will not distinguish between parameter sensitivities and logarithmic gains and will use only the latter term. A logarithmic gain with a magnitude greater than one implies amplification of the original signal; a magnitude less than one indicates attenuation. A positive sign for the logarithmic gain indicates that the changes are in the same direction (both increase, or both decrease in value), while a negative sign indicates that the changes are in the opposite direction. Logarithmic gains can be calculated directly from Eq. 10 as follows:

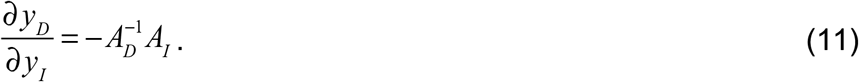

In the context of the Design Space formalism, Eq. 11 implies that the calculation of logarithmic gains does not involve parameter sampling or numerical integration of the system of differential equations. Consequently, logarithmic gains can be used to identify metabolic engineering strategies in a computationally efficient way as illustrated in Section 4.2.

#### 3.1.3 Systems Design Space and Linear Programming

Linear algebra and linear programming play a central role in mathematically defining and characterizing biochemical phenotypes (Fasani and Savageau, 2010). A set of dominant processes, along with an accompanying set of inequalities are only considered to represent a valid phenotype if (a) the S-system equations (Eqs. 3 and 4) have a valid steady-state solution, (b) the set of inequalities (Eqs. 5 and 6) is mathematically consistent, and (c) introducing the solution into the set of inequalities yields a consistent system. Note that step (a) is performed using linear algebra, while steps (b) and (c) involve solving a linear program in each case. Throughout this work, **the Design Space Toolbox v3.0, DST3 (Valderrama-Gómez et al. 2020)**, was used to automatically define and solve linear programs using GLPK as the linear solver. Eqs. S25 to S40 exemplify S-system equations defining phenotype **7306_3**, which will be of interest in later analyses. This set of equations has a valid steady-state solution, which is shown in Eqs. S58 to S73. Associated dominance conditions are represented by Eqs. S41 to S57. Substituting the steady-state solution into the dominance conditions yields Eqs. S74 to S85, which represent phenotypic boundaries delimiting a region in Design Space in which phenotype **7306_3** is valid. A similar process is required to test each one of the potential phenotypes of a given network. DST3 efficiently automates this process, while opening new applications of linear programming in the context of mechanistic modeling, as we will demonstrate in Sections 4.3 and 4.4. For conciseness, mathematical equations defining biochemical phenotypes of the amorphadiene network will not be provided explicitly but can be trivially retrieved using the accompanying Jupyter Notebooks.

### 3.2 The Design Space Toolbox v.3.0 & Jupyter Notebooks

The Design Space Toolbox v.3.0 (DST3) is freely available for all major operating systems through Docker. After Docker has been installed on your system, running the following commands on a terminal window will provide access to DST3:

1. docker pull savageau/dst3**:3.08.79**
2. docker run -d -p 8888:8888 savageau/dst3**:3.08.79**
3. Access the software by opening the address http://localhost:8888/ on any internet browser.

Please refer to the original publication (Valderrama-Gómez et al. 2020) for detailed installation instructions and troubleshooting. Several Jupyter notebooks are provided to reproduce the modeling results of each section. Notebooks can be found within the Docker Image **savageau/dst3:3.08.79** under the directory **/Supporting_Notebooks/AMO_System**. The source code is available under https://github.com/m1vg.

## 4. Results

### 4.1 The Phenotypic Repertoire of the Amorphadiene Biosynthetic Network

We start our analysis by recasting the system of differential equations describing the dynamics of the amorphadiene network (Eqs. S1 to S8) into a fully equivalent GMA form (Eqs. S9 to S24), in which all terms are expressed using power laws. This format is suitable for analysis using DST3 (Valderrama-Gómez et al. 2020). In a following step, the network’s phenotypic repertoire is enumerated along with dynamic properties (number of eigenvalues with positive real part) and volume of individual phenotypes in the System Design Space. This information is summarized in the Supplementary File 1. Note that co-dominant phenotypes were considered for this system (refer to Supplementary Section 4 for details). The amorphadiene network exhibits 40 physiological phenotypes within a parameter range of 10^−3^ to 10^3^ in all dimensions and a total logarithmic volume of 6.25×10^16^. This implies that only about 0.0367% of the 26-dimensional Design Space (with a total logarithmic volume of 6^26^) can support viable biological phenotypes. These numbers suggest that identifying stable operating points by randomly sampling the parameter space is highly inefficient, which is a technique commonly used by ensemble modeling approaches to parameterize mechanistic models (Lee et al., 2014).

The phenotypic repertoire can be filtered to identify top-performing phenotypes without *a priori* knolwedge of parameter values. For that, we use the maximum fold change in production flux from a phenotype’s nominal operating point as the performance metric. When evaluating specific phenotypic properties that depend on a phenotype’s operating point (e.g., eigenvalues, parameter tolerances, phenotype-specific mutation rates (Valderrama-Gómez and Savageau, 2021)), linear programming methods can be applied to identify a nominal parameter set (Lomnitz and Savageau, 2016). The results of this phenotypic assessment are summarized in Fig. S1 as a heatmap. Phenotypes **6921_3, 6913_3, 5769** and **7306_3** exhibit the highest potential to increase production flux from their respective operating point. Phenotype **7306_3** is of central interest in this study, because, as we will show below, it contains the operating point of one of the strains characterized by Weaver et al. (2015).

Available experimental data (such as multi-omics and enzyme kinetics) can be integrated within the phenotype-centric modeling strategy to create a link between observable phenotypic features and regions in Design Space. In the case of the base strain mbis3, experimentally determined protein concentrations (Table S1), along with kinetic parameters extracted from the literature (Table S2), locate the system’s operating point within phenotype **7306_3**, as indicated in Fig. 2A by the black circle. Interestingly, this phenotype was identified in our previous analysis as a potential top performer, which means that the mathematical abstraction for phenotype **7306_3** can be readily used in combination with linear programming to engineer mbis3 into a high performing production strain. Dynamic properties of the strain mbis3 can be determined by an eigenvalue analysis of phenotype **7306_3** (Table S3), which predicts a stable steady state (all eigenvalues are negative real). Numerical integration of the full system of differential equations confirms this prediction (Fig. 2B). Parameter perturbations can move the system’s operating point outside of phenotype **7306_3**. For instance, increasing the concentration of the enzyme amorphadiene synthase (ADS) shifts the operating point from phenotype **7306_3** to **7330_3**, as shown in Fig. 2A by the diamond-shaped symbol. Since the S-system representing phenotype **7330_3** exhibits complex conjugate eigenvalues with positive real part at the denoted operating point (Table S3), the intracellular metabolite concentrations of the network are predicted to exhibit an oscillatory behavior, which is confirmed in Fig. 2C by numerical integration. The ADS concentration can be further increased to place the operating point within a region in Design Space lacking a physiological phenotype (Fig. 2A, black triangle). Thus, some metabolites (MevP, MevPP, and IPP) will not reach a steady state, as shown in Fig. 2D. This behavior arises due to metabolic imbalances present in pathological regions of the Design Space.

**Figure 2.**
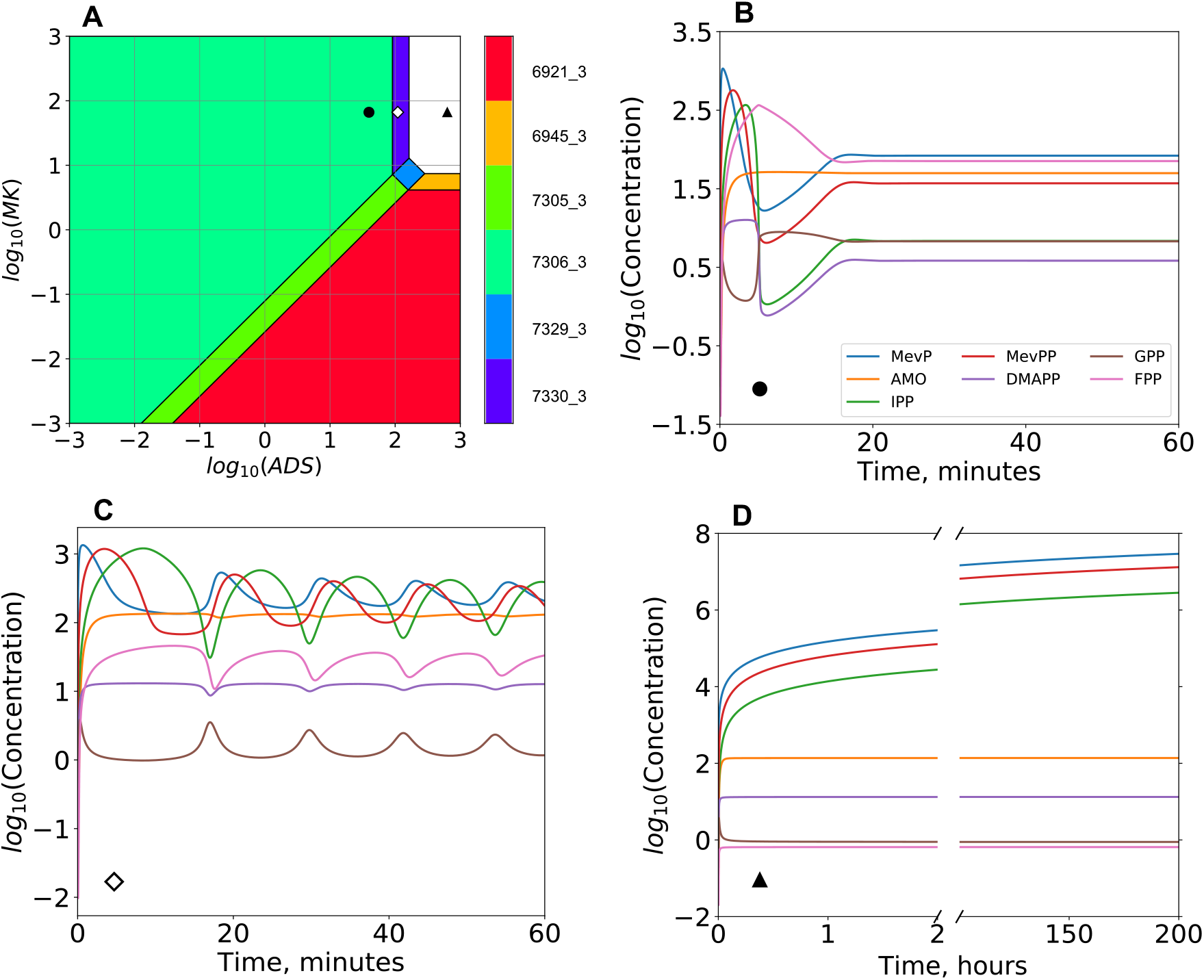
A System Design Space Plot and Three Dynamic Regimes of the Amorphadiene Network. **A**. A Design Space Plot of the amorphadiene biosynthetic network generated for the system defined by Eqs. S9 to S24. Color-coded regions represent biochemical phenotypes of the network. The black circle within phenotype **7306_3** represents the operating point for the base strain mbis3. Kinetic parameters and experimentally determined enzyme concentrations for mbis3 can be found in Tables S1 and S2, which are reproduced from Weaver et. al. (2015). The white diamond symbol within phenotype **7330_3** and the neighboring black triangle in the upper right white region represent hypothetical strains in which the log_10_ concentration of ADS is increased to 2.04 and 2.8, respectively. Panels **B** to **D** show the temporal behavior of intracellular metabolic pools for these three operating points. Each case differs from the other solely by the concentration of ADS. Initial metabolite concentrations were assumed to have a value of 0.1 μM except for mevalonate, whose concentration was set to have a constant value of 5 μM. Numerical integration was performed using the ODEINT routine of SciPy and 10,000 steps. **B**. Stable network dynamics for the base strain mbis3 (black circle in panel **A**). **C**. Oscillatory network dynamics resulting from increasing log_10_(ADS) to 2.04 (white diamond in panel **A**). **D**. Pathological system dynamics when log_10_(ADS) is further increased to 2.8 (black triangle in panel **A**). Note that MevP, MevPP and IPP can no longer reach a steady state but continuously accumulate over time.

The different dynamic regimes shown in Fig 2. can be rationalized in terms of the operation of an integral control system (Aström and Murray, 2010), and the saturation of the enzyme ISPA. The control system, mechanistically implemented by the feedback inhibition of MK by FPP, integrates the difference between the pathway input and output flux to produce the error signal FPP. For example, an increase in ADS increases the output flux and initially decreases its substrate FPP. The decrease in FPP causes de-inhibition of MK and an increase in the input flux until it matches the increased output flux and the change in the error signal FPP goes to zero. As ADS is increased from its initial operating point (Fig. 2B), a switch to phenotype **7330_3** occurs, leaving the integral control system at the boundary of instability and causing oscillations in the concentration of intracellular metabolites (Fig. 2C). With still further increases in ADS and consequent decreases in FPP, there is further de-inhibition of MK to the point that the increase in the input flux exceeds the V_max_ of ISPA, which then becomes the rate limiting step (see second row of Table 1). The saturation of ISPA leads to a new steady state for FPP that is lower with each further increase in ADS, while GPP, DMAPP and AMO maintain a new steady state dictated by the rate limiting flux through ISPA. The de-inhibition of MK and increased input flux causes a buildup of metabolites behind the ISPA bottleneck: MevP, MevPP and IPP (Fig. 2D).

**Table 1.** Enzyme Perturbations Enabling Higher Expression Levels of ADS. Maximal ADS expression values supported by phenotype **7306_1** are listed for multiple conditions. The first row corresponds to the maximal ADS expression supported by the base strain mbis3. Rows 2 to 5 represent hypothetical strains resulting from perturbations of a given set of enzymes (first column) by an amount indicated in the second column.

| Perturbed Enzymes | Fold-Change From mbis3 | Maximal ADS Expression Value ($\mu\text{M}$) |
| --- | --- | --- |
| **ADS** | +2.3 | 91.21 |
| **ISPA, ADS** | +1.79, +4.12 | 163 |
| **PMD, ISPA, ADS** | +1.69, +3.03, +6.97 | 276 |
| **IDI, PMD, ISPA, ADS** | +1.59, +2.69, +4.82, +11.1 | 439 |
| **PMK, IDI, PMD, ISPA, ADS** | +1.91, +3.04, +5.14, +9.2, +21.18 | 838 |

Note that the intervention strategies listed in Table 1 can also be obtained from a manual boundary analysis for phenotype **7306\_3** (Eqs. S74-S85). For instance, the maximal ADS value in the first row of Table 1 is dictated by Eq. S79. The same

### 4.2 A Logarithmic Gain Analysis Reveals a Global Landscape of Metabolic Engineering Strategies

Here, we calculate logarithmic gains in production flux for each one of the 40 physiological phenotypes of the amorphadiene network. The goal is to identify system parameters with the potential to increase amorphadiene productivity. Engineering strategies can be identified from a logarithmic gain analysis using the following rationale: increasing the value of a parameter will enhance productivity when the parameter exhibits a positive logarithmic gain. Conversely, decreasing its value will increase pathway flux when the parameter exhibits a negative logarithmic gain. The global landscape of metabolic engineering strategies based on a logarithmic gain analysis is shown in Fig. 3. It can be divided into four different phenotypic groups according to non-zero logarithmic gains for kinetic parameters associated with a characteristic enzyme set. Each group consists of 10 phenotypes. The first group is characterized by non-zero logarithmic gains for kinetic parameters linked with the last enzyme in the pathway, ADS. The base strain mbis3, whose operating point is located within phenotype **7306_3** (Fig. 2A), belongs to this group. The second and third groups are both characterized by non-zero logarithmic gains for kinetic parameters associated with the first enzyme in the pathway, the mevalonate kinase (MK). Finally, phenotypes within the fourth group exhibit non-zero logarithmic gains for both MK and ADS. Note that Fig. 3 represents the entire landscape of metabolic engineering strategies and is solely based on the architecture of the amorphadiene network (Eqs. S9 to S24). In the context of this analysis, model parameterization is optional and allows the identification of relevant strategies by placing the system’s operating point within one of the four phenotypic groups. In the specific case of strain mbis3, Fig. 3 proposes increasing the concentration of ADS or its turnover number (k_cat7_) as suitable intervention strategies, which is in line with numerical simulations performed by Weaver et al. (2015).

**Figure 3.**
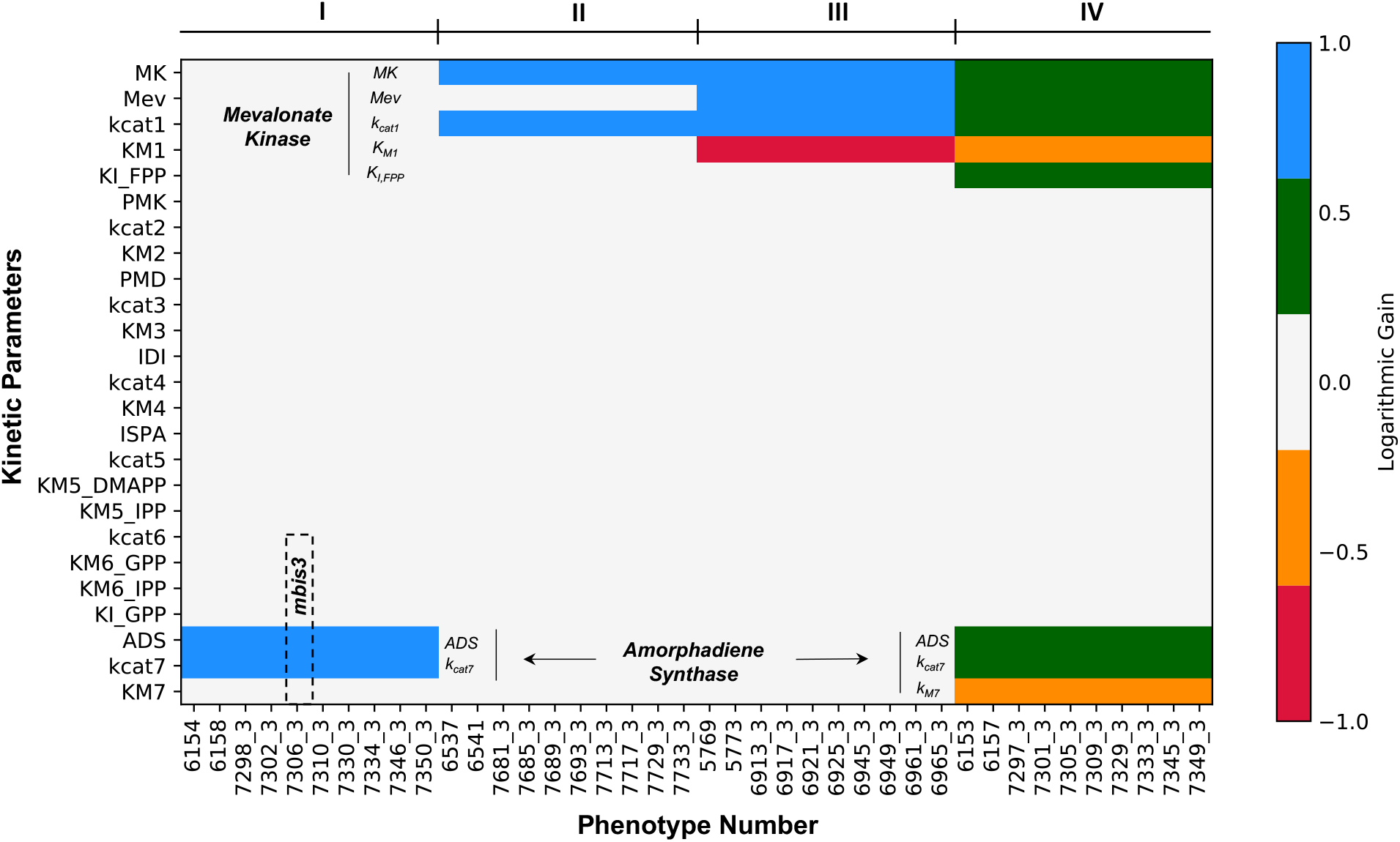
Logarithmic Gains in Pathway Flux Calculated for Physiological Phenotypes. A landscape of 1,040 different logarithmic gains calculated for 40 biochemical phenotypes (x-axis) across 26 system parameters (y-axis) for the amorphadiene production flux, *L(r*_*out*_, *i)*, is shown as a heat map. Logarithmic gain values are color-coded; the white background represents a logarithmic gain of 0. Blue represents a value of 1, green corresponds to 0.5, orange to -0.5 and red corresponds to -1. The rectangle with a black dashed outline highlights phenotype **7306_3**, which contains the operating point of the base strain mbis3. Refer to Supplementary File 3 for the figure’s raw data.

Phenotype signatures (Savageau et al. 2009, Valderrama-Gómez et al. 2020) encode necessary instructions to construct S-system equations (i.e., Eqs. 3 and 4) from the full system of differential equations. From a biological point of view, these signatures contain information about dominant fluxes and biochemical mechanisms exercised by each phenotype. An analysis of *conserved* dominance signatures (Table S4 and S5) for individual groups in Fig. 3 reveals that saturation patterns of two key enzymes (MK, and ADS, see Table. S6 and Fig. S2) can be used to differentiate individual phenotypic groups. Further, this suggests that a targeted enzymatic characterization has the potential to rapidly assign a given strain to one of these four groups. For instance, groups II and III differ solely by the saturation features of MK (Fig. S2). Since the mevalonate concentration is the largest positive term in Eq. S16 for all phenotypes within group II, we consider the enzyme MK to be saturated. Conversely, the same enzyme is *not* saturated in group III because K_M1_ is the largest positive term in Eq. S16. These saturation differences directly influence engineering strategies, as evidenced in Fig. 3 (compare groups II and III). When MK is saturated, increasing the mevalonate kinase concentration (MK) or its turnover number (k_cat1_) are the only two possible strategies to significantly increase amorphadiene productivity. However, if MK is *not* saturated, increasing the mevalonate concentration and decreasing the Michaelis-Menten constant K_M1_ are two additional strategies that can be implemented to increase productivity. Several other analogous comparisons can be made. Let us consider the saturation patterns of phenotypic groups III and IV. At a first glance, there are no obvious differences in the enzyme saturation pattern of these two groups (see Table S6 and Figure S2). In each case, the key enzymes MK and ADS are not saturated. However, a detailed inspection of the MK saturation regime in group IV reveals an important difference. While K_M1_ is the largest positive term in Eq. S16 for all phenotypes within group III, the inhibition term K_M1_ * FPP * K^-1^_I,FPP_ is the largest one in group IV. The consequences of this subtle difference in dominance are reflected in additional engineering strategies involving not only MK but also ADS in group IV (refer to Fig. 3, groups III vs. IV). Increasing K_I,FPP_ positively impacts productivity in group IV because it reduces the aggregate Michaelis-Menten constant for MK (K_M1_ * FPP * K^-1^_I,FPP_ + K_M1_). Additionally, increasing ADS, k_cat7_ or decreasing K_M7_ accomplishes the same goal by decreasing the steady state value of the metabolic intermediate FPP (Fig. 1) –which can be inferred by a logarithmic gain analysis for the exemplary phenotype **7333_3** within group IV: L(FPP, ADS) = -0.5, L(FPP, k_cat7_)= -0.5, and L(FPP,K_M7_)= 0.5.

### 4.3 Identifying Coupled Enzyme Targets

Using numerical simulation and a partial rank correlation analysis for a set of 10,000 models, Weaver et al. (2015) identified the concentration of ADS and its turnover number (k_cat7_) as the most relevant parameters to increase amorphadiene productivity. To refine model predictions, the authors experimentally determined the *in vivo* value for k_cat7,_ which they estimated to be 0.022 ± 0.008 s^-1^. This number is significantly different from the value extracted from the literature (0.0068 s^-1^). Fig. 4A shows a Design Space plot for the amorphadiene network with an updated k_cat7_ value. Note that the qualitative arrangement of neighboring phenotypes, as well as the location of the operating point of the base strain mbis3 within phenotype **7306_3**, remain unchanged when the k_cat7_ value extracted from the literature is used instead (refer to the Supplementary IPython Notebook 4.1 within the DST3 Docker Image).

**Figure 4.**
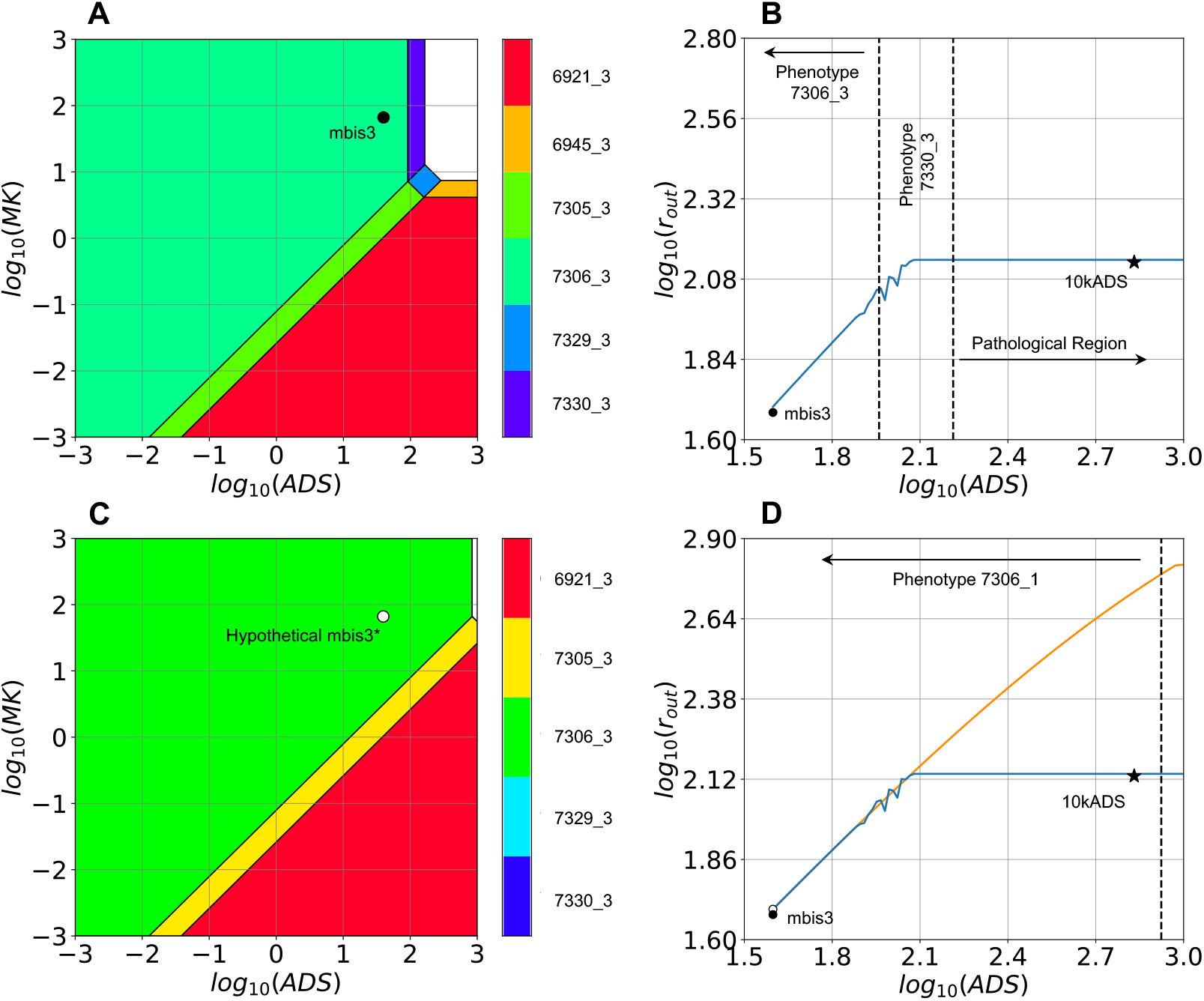
System Design Space Plots and *in silico* Titration Studies. A. A Design Space plot for strain mbis3 (black circle within phenotype **7306_3**) is shown. **B**. A titration plot (solid blue line) is generated for the ODE system (Eqs. S1 – S8) by increasing the expression level of ADS from its basal concentration of 39.6 μM in strain mbis3 to 1,000 μM. The two vertical dashed lines mark the boundaries of phenotype **7330_3**. Operating points located within this biochemical phenotype have the potential to exhibit oscillatory behavior, qualitatively similar to the one shown in Fig. 2C. The average slope of the titration curve within the boundaries of phenotype **7306_3** is 0.98, which closely matches a logarithmic gain of L(r_out_, ADS) = 1 calculated for phenotype **7306_3** (Fig. 3) using linear algebra. Experimental amorphadiene production rates for strains mbis3 and 10kADS are represented by a black star and a black circle, respectively. **C**. A Design Space plot is shown for the hypothetical strain mbis3* (white circle within phenotype **7306_3**). This strain results from increasing the levels of PMK (1.91-fold), IDI (3.04-fold), PMD (5.14-fold), and ISPA (9.2-fold), from the respective levels of the base strain mbis3. **D**. A titration plot for the ODE system (solid orange line) is generated by increasing the expression level of ADS from its basal concentration of 39.6 μM in the hypothetical strain mbis3* (white circle) to 1,000 μM. The blue solid line corresponds to the titration study performed in panel **B**. The vertical dashed line represents the right boundary of phenotype **7306_3**. Experimental productivity of the strains mbis3 and 10kADS are represented by a black circle and a black star, respectively. The average slope of the titration curve within phenotype **7306_1** is 0.82, in close agreement with a logarithmic gain of L(r_out_, ADS) = 1 calculated for the same phenotype. Average slopes are determined by computing two-point slopes and averaging their values over the entire curve.

To test the effect of increasing ADS expression on amorphadiene productivity, Weaver et al. (2015) constructed and characterized the strain 10kADS, which contains a stronger ribosome binding site in front of the ADS sequence. The experimentally determined ADS concentration in 10kADS was 678 μM, which corresponds to a 17-fold increase compared with its level in the base strain mbis3 (39.6 μM). As reported by the authors, there is a good agreement between model predictions (blue line in Fig. 4B) and the experimental performance of strain 10kADS (black star in Fig. 4B). Our logarithmic gain analysis for phenotype **7306_3** not only predicts the fact that increasing ADS expression will increase amorphadiene productivity (Fig. 3), but it also provides an estimate for the magnitude of such an increase. A good agreement is observed between the slope of the blue curve to the left of the leftmost vertical line in Fig. 4B, whose value corresponds to 0.98, and the calculated logarithmic gain for the pathway flux, L(r_out_, ADS) = 1, for phenotype **7306_3**. Note that an experimental log_10_(ADS) value of 2.8 places the operating point for strain 10kADS outside the boundaries of phenotype **7306_3**, well within the pathological region to the right of phenotype **7330_3** (Fig. 4A and B). The location of the operating point of strain 10kADS is relevant because the predicted positive effect of ADS overexpression on amorphadiene productivity is only valid within the boundaries of phenotype **7306_3**, for which a logarithmic gain of L(r_out_, ADS) = 1 is calculated. As shown in Fig. 4B, after log_10_(ADS) surpasses the right boundary of phenotype **7330_3** (rightmost vertical dashed line), further increasing ADS does not translate into a higher amorphadiene synthesis rate. Instead, overexpressing ADS at higher levels can potentially decrease strain performance due to the accumulation of toxic metabolites (MevPP, MevP and IPP, see Fig. 2D) and the associated protein burden.

Since L(r_out_, ADS) = 1 is only valid within the boundaries of phenotype **7306_3** (Eqs. S74-S85), identifying additional engineering targets to increase pathway performance is analogous to identifying enzyme perturbations that allow higher ADS expression levels within the boundaries of phenotype **7306_3**. We exploit linearities in the mathematical definition of biochemical phenotypes in logarithmic space to formulate and solve this optimization task using linear programming. Starting from the operating point of the base strain mbis3, we allow the concentration of a set of enzymes, including ADS, to vary within the range 10^−3^ to 10^3^. Note that this range can be easily adjusted if needed. Then, the concentration of the free enzymes is adjusted so that the expression of ADS is maximal within the boundaries of phenotype **7306_3**. We perform this procedure varying the number of free enzymes from 1 to 4. The results are summarized in Table 1. Adjusting the expression level of the enzymes PMK, IDI, PMD, and ISPA as indicated in the last row of Table 1, would allow the resulting hypothetical strain mbis3* (Fig. 4C) to support a maximal ADS expression of 838 μM, which is higher than the ADS level in strain 10kADS (678 μM). The effect of increasing ADS to this level on amorphadiene productivity is shown in Fig. 4D by the intersection of the orange curve and the dashed vertical line. As ADS is increased, so does the production flux through the amorphadiene network. The average slope of the orange curve to the left of the vertical dashed line is 0.82 and agrees well with a predicted logarithmic gain for phenotype **7306_3** of L(r_out_, ADS)=1.0. The net effect of adjusting the concentrations of PMK, IDI, PMD and ISPA is to extend the region of validity of phenotype **7306_3** (compare Figs. 4A and 4C). Consequently, the network could support a productivity of 608 μM/min, which represents a 4.5-fold increase compared with the experimental productivity for strain 10kADS (135 μM/min or 2.25 μM/s).

Note that the intervention strategies listed in Table 1 can also be obtained from a manual boundary analysis for phenotype **7306_3** (Eqs. S74-S85). For instance, the maximal ADS value in the first row of Table 1 is dictated by Eq. S79. The same equation also points to ISPA as the first enzyme whose level needs to be fine-tuned to allow for a higher ADS expression. Furthermore, Eq. S78 identifies PMD as an additional target, followed by IDI and PMK, which are identified through Eqs. S74 and S77, respectively. An automated analysis of phenotypic boundaries by linear programming will be the method of choice as the network’s scope and number of inequalities increase.

Since experimentally implementing the fold-change values listed in Table 1 for perturbed enzymes might be challenging because of technical difficulties associated with continuous titration of enzyme levels, *integer* linear programming (Schrijver, 1986) provides an alternative approach to increase the biological feasibility of the strategies identified from a boundary analysis for phenotype **7306_3**. From an experimental point of view, these intervention strategies could be implemented more easily, for instance, by fine-tuning the copy number of plasmids harboring target enzymes. The biological feasibility of the predictions could be further refined by considering appropriate constraints on the total number of plasmids that can be supported by the cell.

### 4.4 Modulating Feedback Inhibition as a Valid Engineering Strategy

Mevalonate kinase (MK), the first enzyme of the pathway, is subjected to feedback inhibition by the pathway intermediate farnesyl diphosphate (FPP) (Miziorko, 2011). Weaver et al. (2015) hypothesized that alleviating feedback inhibition in MK could increase amorphadiene productivity. A logarithmic gain analysis for biochemical phenotype **7306_3** (Fig. 3), which contains the operating point of the base strain mbis3 (Fig. 4A), indicates that the pathway flux (r_out_) is *insensitive* to perturbations in K_I, FPP_, i.e., L(r_out_, K_I,FPP_) = 0.0. Similarly, but using numerical simulation, Weaver et al. (2015) observed no significant effect of perturbations in K_I,FPP_ on pathway flux. To experimentally validate this observation, the authors constructed and characterized strain saMK containing a homologous mevalonate kinase from *Staphylococcus aureus* with a 24-fold weaker K_I, FPP_. In line with our logarithmic gain analysis and numerical simulations by Weaver et al. (2015), the experimental amorphadiene production in strain saMK was not sensitive to mevalonate kinase inhibition.

We now employ the phenotype-centric approach to answer two related questions: a) does a biological phenotype exist for which modulating the MK feedback inhibition is a valid strategy to increase amorphadiene productivity? and b) what is the effect of completely removing MK inhibition on the global landscape of valid engineering strategies increasing amorphadiene production? We use Fig. 3 to answer the first question. A logarithmic gain of L(r_out_, K_I,FPP_) = 0.5 in the fourth phenotypic group points to the existence of ten different biochemical phenotypes harboring operating points for which increasing values of K_I,FPP_ lead to an increased amorphadiene productivity. These phenotypes are listed in Table 2, along with their phenotypic volumes in logarithmic space (a proxy of phenotypic robustness) and non-zero logarithmic gains for pathway flux. To demonstrate the power of the phenotype-centric strategy to efficiently explore the Design Space using linear programming, we aim to identify an operating point within phenotype **6153** fulfilling two conditions: K_I,FPP_ > 1.9 μM and r_out_ > 50 μM/min. These constraints are employed to identify an operating point which is comparable with that of the base strain mbis3. Starting from the mbis3 operating point, parameter values of a “free set” (which includes K_I, FPP_) are allowed to vary within the range 10^−3^ to 10^3^. If phenotype **6153** is valid within the resulting high-dimensional cube, the tolerance for K_I,FPP_ is calculated (minimum and maximum value) using linear programming. This procedure was initially performed for free sets containing only protein concentrations (MK, PMK, PMD, IDI, ISPA and ADS). The underlying idea was to identify an operating point within phenotype **6153** that could be experimentally reached starting from mbis3 by simply adjusting the expression of a given set of enzymes. However, this was not possible for free sets of any size (n=1, 2, …, 5 and 6). Thus, free sets were expanded to consider not only protein concentrations, but also enzyme kinetic parameters. Fig. 5A shows the location of one of such operating points (white circle) within phenotype **6153** obtained using this procedure. We term this operating point mbis3**. Reaching mbis3** requires fine tuning ADS, ISPA, and K_M7_ to have values of 163 μM, 105.9 μM and 1,000 μM, respectively, while keeping all other kinetic parameters of the base strain mbis3 unchanged. As shown in Fig. 5B, alleviating MK feedback inhibition in the hypothetical strain mbis3** leads to a 4-fold improvement in productivity. This is in stark contrast with the experimental performance of strain saMK, whose amorphadiene productivity remained almost unchanged after alleviating MK inhibition by the same extent.

**Table 2.**
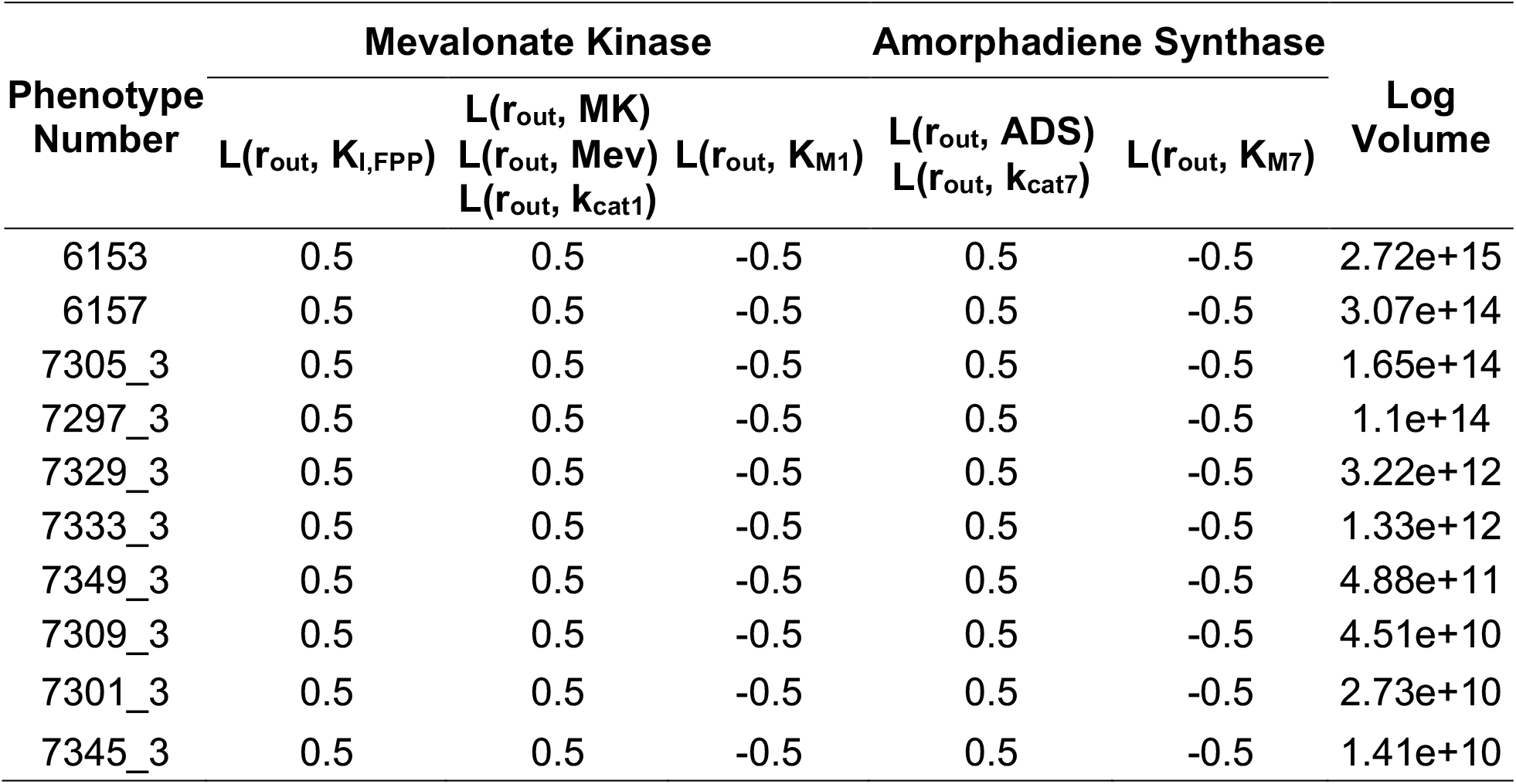
Biochemical Phenotypes Responding to Changes in K_I,FPP_. Relevant logarithmic gains and phenotypic volumes in logarithmic space are listed for ten phenotypes for which L(r_out_, K_I,FPP_) ≠ 0. In each case, the pathway flux (r_out_) can be increased by modifying parameter values associated with either MK or ADS. Phenotypic volumes are calculating using the tolerance method, which provides an underestimate (Valderrama-Gómez and Savageau, 2021). By virtue of its volume, phenotype **6153** is considered as the most robust to parametric perturbations.

**Figure 5.**
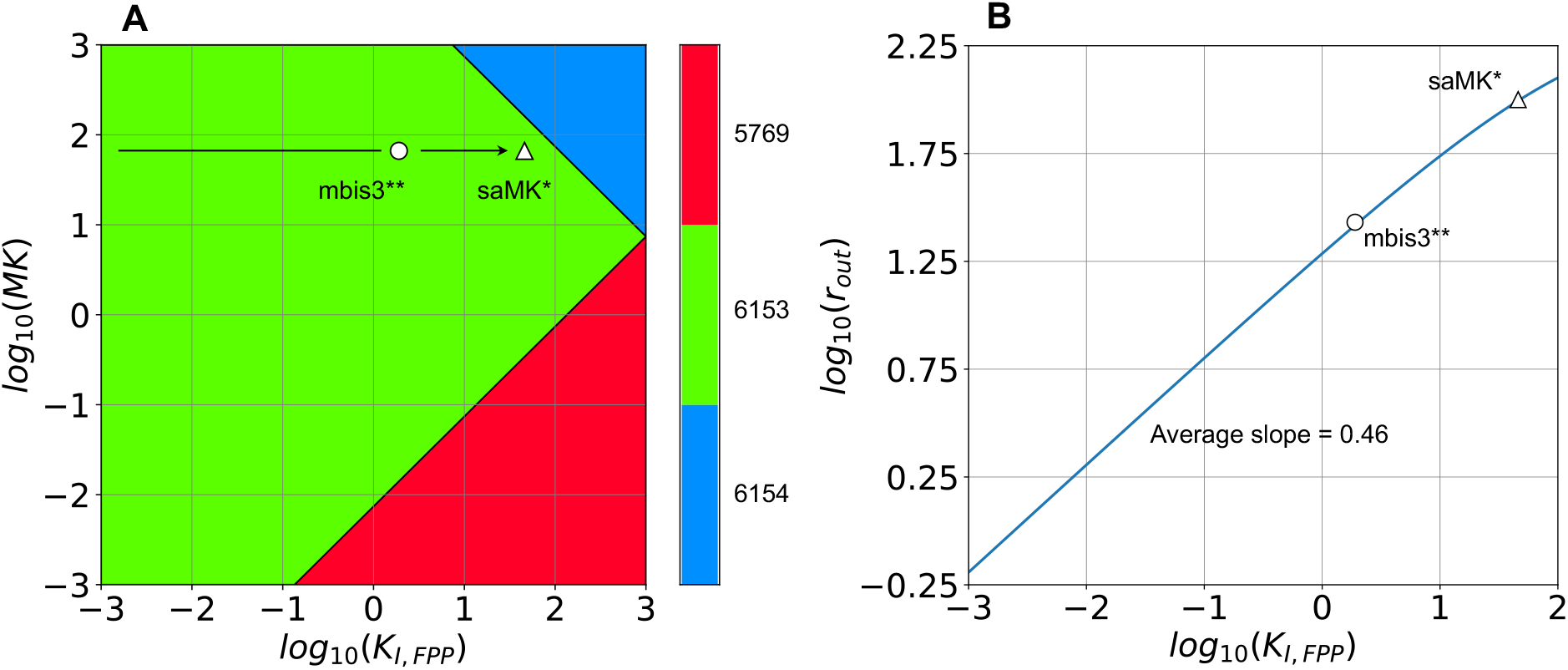
Design Space and Titration Plots for Phenotype 6153. **A**. The relative locations of strains mbis3** (white circle) and saMK** (white triangle) within phenotype **6153** are shown. Mbis3** is a hypothetical strain resulting from setting ADS = 163 μM, ISPA = 106 μM and K_M7_ = 1,000 μM while keeping all other kinetic parameters of the base strain mbis3 unchanged. Strain saMK* is also a hypothetical strain that results from alleviating MK inhibition in mbis3** by increasing its K_I,FPP_ from 1.9 μM to 46 μM. **B**. The estimated amorphadiene productivity for mbis3** corresponds to 27 μM/min and is lower than the base strain mbis3, which corresponds to roughly 48 μM/min (Fig. 5 in Weaver et al., 2015). The predicted amorphadiene productivity of strain saMK* is 100 μM/min, which is over 2.5-fold higher than the experimentally determined value for the strain saMK (Fig. 5 in Weaver et al., 2015). The slope of the titration plot agrees well with a predicted logarithmic gain of L(r_out_, K_I,FPP_) = 0.5 for phenotype **6153**.

We now turn our attention to the second question: what is the effect of completely removing MK inhibition on the global landscape of valid engineering strategies increasing amorphadiene production? As demonstrated in Section 4.2, the phenotype-centric approach can be used to elucidate the mechanistic link between a network’s architecture and its function. Thus, we explore *in silico* the structural effect of completely removing MK feedback inhibition (Fig. S5A) on engineering strategies increasing production flux. As shown in Fig. S5B, removing K_I,FPP_ from Eqs. S16 eliminates phenotypic groups I and IV from the original landscape of metabolic engineering strategies (Fig. 3). Interestingly, and as a direct consequence of this structural modification, a logarithmic gain analysis suggests that overexpressing ADS or increasing its turnover number k_cat7_ would no longer increase pathway flux, as was the case for strain mbis3 (located within phenotypic group I). A simple mathematical analysis (refer to Supplementary Section 5) can be used to calculate equivalent operating points for the modified network without feedback inhibition. Figs. S5C and S5D show the location of two such operating points. Since in each case the operating points are located on a phenotypic boundary involving MK, overexpressing the concentration of this enzyme to increase amorphadiene productivity can potentially lead to metabolic imbalances and a decreased product yield.

## 5. Discussion

The notion that predictions from kinetic models are context specific and only informative under the conditions for which the underlying kinetic model has been parameterized is commonly accepted (Chowdhury et al., 2015). However, to the best of our knowledge, a mathematical formalism that generates predictions while providing the biological context in which those predictions are valid is still missing in the field of rational Metabolic Engineering. In Section 4.2, we showed that the phenotype-centric strategy generates predictions from mechanistic models without requiring parameterization or numerical integration of the underlying system of differential equations. Additionally, we showed that the context in which model predictions are valid is provided by the boundaries of the biochemical phenotype from which those predictions stem (Fig. 4B). For the case study analyzed here, Weaver et al. (2015) increased amorphadiene productivity by overexpressing the enzyme ADS from a low level in the base strain mbis3 to a high level in the production strain 10kADS. Since the biological context in which the positive effect of ADS overexpression on productivity was not considered to fine-tune ADS expression experimentally, a too high ADS level in strain 10kADS effectively placed its operating point within a pathological region of the Design Space (Fig. 4B). Consequently, metabolic intermediates (MevP, MevPP and IPP) have the potential to accumulate to toxic levels (Fig. 2D) in this strain. By confining ADS overexpression to the boundaries of the physiological phenotype **7306_3**, as shown in Fig. 4B, the resulting production strain could have the potential to outperform strain 10kADS in terms of product yield, due to balanced intracellular metabolite levels and a reduced protein burden. Strains mbis3^a^ and 10kADS^b^ in Fig. S3 illustrate this point.

The role of feedback inhibition is critical to the operation of the amorphadiene network. It is essential for the implementation of the integral control that matches input flux to the output flux that is determined by the saturation of ADS. Elimination of this feedback inhibition can cause one of three behaviors, depending on the uncontrolled rate of the mutated MK: a) If the uncontrolled rate of MK is greater than that of ADS, there will be a continual increase of material withing the pathway (a pathological phenotype characterized by a blowup), b) if the uncontrolled rate of MK is less than that of ADS, there will be a continual decrease of material withing the pathway (a “blowdown”). However, in this case ADS will eventually become unsaturated, and the pathway will come to a new steady state and c) if the uncontrolled rate of MK is equal to that of ADS, as described in the previous section and Supplementary Section 5, there will be a steady state that is marginally stable and any transient reduction of metabolites from the pathway will lead to another marginally stable steady state with less material being held within the system. There is no unique steady state solution but rather a marginally stable manifold of solutions. The detailed analysis of a simplified system that clearly exhibits this tripartite behavior can be found in the appendix of Savageau (1969).

The phenotype-centric modeling strategy enables a computationally efficient exploration of the System Design Space at two different levels of detail. The first one is conducted at a global scale, involves the enumeration of the system’s phenotypic repertoire (Section 4.1) and the semi-quantitative characterization of phenotypic properties, such as robustness, dynamic behavior, and logarithmic gains (Section 4.2). This step is automatically performed by the Design Space Toolbox v.3.0 (Valderrama-Gómez et al., 2020) and does not require *a priori* knowledge of kinetic parameters. In contrast, the second level involves specific numerical values for the system’s parameters and is relevant when identifying a robust operating point within a biochemical phenotype of interest (Valderrama-Gómez et al., 2020), maximizing the region of validity for a specific phenotypic trait (Section 4.3), or identifying efficient transitions in parameter space between biochemical phenotypes (Section 4.4). At this level of detail, ensemble modeling approaches addressing parametric uncertainties by dense sampling (Tran et al., 2008; Lee et al., 2014) could benefit from the ability of the phenotype-centric modeling strategy to identify physiological phenotypes. This synergy would dramatically speedup model parameterization by restricting parameter sampling to regions in Design Space leading to stable, physiological models. Further, conventional optimization methods, such as gradient descent (Ruder, 2017), Newton’s method (Polyak, 2007), and evolutionary algorithms (Bäck and Schwefel, 1993), among others, could also benefit from an efficient identification of regions of interest in Design Space exhibiting desired properties.

Regardless of the level in which the Design Space is explored, numerical simulation of the underlying system of differential equations (e.g., Figs. 2B-D, 4B, 4D and 5B) is not required in the context of the phenotype-centric modeling strategy and is only performed in this work to confirm our predictions. Overall, we observed a high accuracy in our predictions, as evidenced by (a) logarithmic gains estimated for biochemical phenotypes **7306_3** (Fig. 4B and D) and **6153** (Fig. 5B) closely matching the slopes of the respective titration curves, and (b) successful prediction of the full system’s dynamics using an eigenvalue analysis of relevant biochemical phenotypes (Fig. 2 and Table S3). Note that deviations in our predictions from the actual behavior of the full system are a natural consequence of the mathematical definition of biochemical phenotypes. Deviations are expected to be low about a phenotype’s centroid and higher at phenotypic boundaries, where, by definition, there is no dominance (Savageau and Lomnitz, 2014).

Even though the mechanistic model analyzed here only considered enzyme-catalyzed metabolic processes, the mathematical formalism behind the phenotype-centric approach is general and can handle models covering protein and mRNA synthesis with multiple regulatory layers at the transcription, translation and post-translation levels. The only formal requirement is that the mechanisms are described by fundamental chemical and biochemical kinetics, which can be recast into the GMA form as exemplified in the Supplementary Section 2. One additional aspect to consider when building and analyzing kinetic models for Metabolic Engineering applications is the effect of enzyme overexpression on proteome allocation. This effect will become particularly important when one enzyme makes up a significant percentage of the overall proteome due to enzyme overexpression. Note that the effect of ADS overexpression on pathway enzyme levels was not considered in the titration studies shown in Figs. 4B and 4D. However, this can be done by constraining the total concentration of pathway enzymes (or in general, the total proteome) to a given value. We recently expanded the Design Space Toolbox (Valderrama-Gómez et al., 2020) to handle the system of algebraic differential equations resulting from such considerations and we expect to explore the effect of proteome allocation constraints for kinetic models in a future work.

We believe that the phenotype-centric strategy has the potential to advance the field of rational Metabolic Engineering by (a) providing an efficient way to explore the Design Space at different levels of detail, (b) allowing the evaluation of model hypothesis in a structured manner, (c) enabling metabolic network optimization based on kinetic models without requiring *a priori* knowledge of parameter values, and (d) serving as a scaffold for the development of kinetics-based algorithms for rational Metabolic Engineering. Using the amorphadiene biosynthetic network as a case study, we demonstrated each one of these advantages and provided a mechanistic context for the experimental work of Weaver et al. (2015). We envision next generation development of strain-design algorithms and methods for rational pathway optimization to exploit the predictive power of mechanistic models by leveraging a modeling paradigm that is more focused on biochemical phenotypes and their transitions and relies less on first requiring specific parameter values and numerical simulation.

## 6. Acknowledgements

This work was supported in part by a grant from the US National Science Foundation Grant number MCB 1716833.

## 7. Author contributions

Conceptualization, M.A.V.; Methodology, M.A.V. and M.A.S.; Software, M.A.V; Validation, M.A.V.; Investigation, M.A.V. and M.A.S.; Writing, M.A.V. and M.A.S; Funding Acquisition, M.A.S.

