## Supplementary Information for "Phenotype-Centric Modeling for Rational Metabolic Engineering"

### 1. Kinetic Model of Amorphadiene Synthesis from Mevalonate

The model is comprised of seven differential equations (Eq. S1 to Eq. S7) and one algebraic constraint (Eq. S8) and involves 26 kinetic parameters. This model considers an output flux for amorphadiene ($r_{out}$, Eq. S7 and S8) and thus slightly differs from the original model by Weaver et al. (2015), which does not consider this term.

$$\frac{dMevP}{dt} = \frac{k_{cat1}[MK][Mev]}{K_{M1} + K_{M1}[FPP]K_{I,FPP}^{-1} + [Mev]} - \frac{k_{cat2}[PMK][MevP]}{K_{M2} + [MevP]} \tag{S1}$$

$$\frac{dMevPP}{dt} = \frac{k_{cat2}[PMK][MevP]}{K_{M2} + [MevP]} - \frac{k_{cat3}[PMD][MevPP]}{K_{M3} + [MevPP]} \tag{S2}$$

$$\begin{aligned}\frac{dIPP}{dt} = {} & \frac{k_{cat3}[PMD][MevPP]}{K_{M3} + [MevPP]} + \frac{k_{cat4}[IDI][DMAPP]}{K_{M4} + [DMAPP]} - \frac{k_{cat4}[IDI][IPP]}{K_{M4} + [IPP]} \\ & - \frac{k_{cat5}[ISPA][IPP][DMAPP]}{K_{M5,IPP}[DMAPP] + K_{M5,DMAPP}[IPP] + [DMAPP][IPP]} \\ & - \frac{k_{cat6}[ISPA][IPP][GPP]}{K_{I,GPP}K_{M6,IPP} + K_{M6,IPP}[GPP] + K_{M6,IPP}[IPP] + [GPP][IPP]}\end{aligned} \tag{S3}$$

$$\begin{aligned}\frac{dDMAPP}{dt} = {} & \frac{k_{cat4}[IDI][IPP]}{K_{M4} + [IPP]} - \frac{k_{cat4}[IDI][DMAPP]}{K_{M4} + [DMAPP]} \\ & - \frac{k_{cat5}[ISPA][IPP][DMAPP]}{K_{M5,IPP}[DMAPP] + K_{M5,DMAPP}[IPP] + [DMAPP][IPP]}\end{aligned} \tag{S4}$$

$$\begin{aligned}\frac{dGPP}{dt} = {} & \frac{k_{cat5}[ISPA][IPP][DMAPP]}{K_{M5,IPP}[DMAPP] + K_{M5,DMAPP}[IPP] + [DMAPP][IPP]} \\ & - \frac{k_{cat6}[ISPA][IPP][GPP]}{K_{I,GPP}K_{M6,IPP} + K_{M6,IPP}[GPP] + K_{M6,IPP}[IPP] + [GPP][IPP]}\end{aligned} \tag{S5}$$

$$\begin{aligned}\frac{dFPP}{dt} = {} & \frac{k_{cat6}[ISPA][IPP][GPP]}{K_{I,GPP}K_{M6,IPP} + K_{M6,IPP}[GPP] + K_{M6,GPP}[IPP] + [GPP][IPP]} \\ & - \frac{k_{cat7}[ADS][FPP]}{K_{M7} + [FPP]}\end{aligned} \tag{S6}$$

$$\frac{dAMO}{dt}=\frac{k_{cat7}[ADS][FPP]}{K_{M7}+[FPP]}-r_{out} \quad \text{(S7)}$$

$$r_{out}=k_{out}[AMO] \quad \text{(S8)}$$

### 2. Kinetic Model in Generalized Mass Action (GMA) Form

Equations S9 to S24 represent the mechanistic model in a fully equivalent format that involves only sums of products of power laws and is thus suitable for analysis using the Design Space Toolbox v3.0 (Valderrama-Gómez et al. 2020).

$$\frac{dMevP}{dt}=k_{cat1}[MK][Mev]D_1^{-1}-k_{cat2}[PMK][MevP]D_2^{-1} \quad \text{(S9)}$$

$$\frac{dMevPP}{dt}=k_{cat2}[PMK][MevP]D_2^{-1}-k_{cat3}[PMD][MevPP]D_3^{-1} \quad \text{(S10)}$$

$$\begin{aligned}\frac{dIPP}{dt}&=k_{cat3}[PMD][MevPP]D_3^{-1}+k_{cat4}[IDI][DMAPP]D_{4,r}^{-1}\\&-k_{cat4}[IDI][IPP]D_{4,f}^{-1}-k_{cat5}[ISPA][IPP][DMAPP]D_5^{-1}\\&-k_{cat6}[ISPA][IPP][GPP]D_6^{-1}\end{aligned} \quad \text{(S11)}$$

$$\begin{aligned}\frac{dDMAPP}{dt}&=k_{cat4}[IDI][IPP]D_{4,f}^{-1}-k_{cat4}[IDI][DMAPP]D_{4,r}^{-1}\\&-k_{cat5}[ISPA][IPP][DMAPP]D_5^{-1}\end{aligned} \quad \text{(S12)}$$

$$\frac{dGPP}{dt}=k_{cat5}[ISPA][IPP][DMAPP]D_5^{-1}-k_{cat6}[ISPA][IPP][GPP]D_6^{-1} \quad \text{(S13)}$$

$$\frac{dFPP}{dt}=k_{cat6}[ISPA][IPP][GPP]D_6^{-1}-k_{cat7}[ADS][FPP] \quad \text{(S14)}$$

$$\frac{dAMO}{dt}=k_{cat7}[ADS][FPP]D_7^{-1}-r_{out} \quad \text{(S15)}$$

$$0=K_{M1}+K_{M1}[FPP]K_{I,FPP}^{-1}+[Mev]-D_1 \quad \text{(S16)}$$

$$0=K_{M2}+[MevP]-D_2 \quad \text{(S17)}$$

$$0 = K_{M3} + [MevPP] - D_3 \quad \text{(S18)}$$

$$0 = K_{M4} + [IPP] - D_{4,f} \quad \text{(S19)}$$

$$0 = K_{M4} + [DMAPP] - D_{4,b} \quad \text{(S20)}$$

$$0 = K_{M5,IPP}[DMAPP] + K_{M5,DMAPP}[IPP] + [DMAPP][IPP] - D_5 \quad \text{(S21)}$$

$$0 = K_{I,GPP}K_{M6,IPP} + K_{M6,IPP}[GPP] + K_{M6,IPP}[IPP] + [GPP][IPP] - D_6 \quad \text{(S22)}$$

$$0 = K_{M7} + [FPP] - D_7 \quad \text{(S23)}$$

$$0 = k_{out}[AMO] - r_{out} \quad \text{(S24)}$$

### 3. Mathematical Description of Phenotype 7306_3

*S-System Equations*

$$\frac{dMevP}{dt} = k_{cat1}[MK][Mev]D_1^{-1} - k_{cat2}[MevP][PMK]D_2^{-1} \quad \text{(S25)}$$

$$\frac{dMevPP}{dt} = k_{cat2}[PMK][MevP]D_2^{-1} - k_{cat3}[MevPP][PMD]D_3^{-1} \quad \text{(S26)}$$

$$\frac{dIPP}{dt} = k_{cat4}[DMAPP][IDI]D_{4r}^{-1} - k_{cat4}[IPP][IDI]D_{4f}^{-1} \quad \text{(S27)}$$

$$\frac{dDMAPP}{dt} = \frac{1}{3}k_{cat3}[Mev][PPD]D_3^{-1} - k_{cat5}[IPP][DMAPP][ISPA]D_5^{-1}$$

(S28)

$$\frac{dGPP}{dt} = k_{cat5}[IPP][DMAPP][ISPA]D_5^{-1} - k_{cat6}[IPP][GPP][ISPA]D_6^{-1} \quad \text{(S29)}$$

$$\frac{dFPP}{dt} = k_{cat6}[IPP][GPP][ISPA]D_6^{-1} - k_{cat7}[FPP][ADS]D_7^{-1} \quad \text{(S30)}$$

$$\frac{dAMO}{dt} = k_{cat7}[FPP][ADS]D_7^{-1} - k_{out}[AMO] \quad \text{(S31)}$$

$$0 = K_{M1}K_{1,FPP}^{-1}[FPP] - D_1 \quad \text{(S32)}$$

$$0 = K_{M2} - D_2 \quad \text{(S33)}$$

$$0 = K_{M3} - D_3 \quad \text{(S34)}$$

$$0 = K_{M4} - D_{4,f} \quad \text{(S35)}$$

$$0 = K_{M4} - D_{4,r} \quad \text{(S36)}$$

$$0 = [IPP]K_{M5,DMAPP} - D_5 \quad \text{(S37)}$$

$$0 = K_{M6,IPP}K_{I,GPP} - D_6 \quad \text{(S38)}$$

$$0 = FPP - D_7 \quad \text{(S39)}$$

$$0 = k_{out}[AMO] - r_{out} \quad \text{(S40)}$$

*__Dominance Conditions__*

$$2k_{cat5}k_{cat6}^{-1}[DMAPP][GPP]^{-1}D_5^{-1}D_6 > 1 \quad \text{(S41)}$$

$$3k_{cat3}^{-1}k_{cat4}[PMD]^{-1}[MevPP]^{-1}[IDI][DMAPP]D_{4,r}^{-1}D_3 > 1 \quad \text{(S42)}$$

$$k_{cat5}^{-1}k_{cat4}[ISPA]^{-1}[DMAPP]^{-1}[IDI]D_{4,f}^{-1}D_5 > 1 \quad \text{(S43)}$$

$$k_{cat6}^{-1}k_{cat4}[ISPA]^{-1}[GPP]^{-1}[IDI]D_{4,f}^{-1}D_6 > 1 \quad \text{(S44)}$$

$$k_{cat5}^{-1}k_{cat4}[IPP]^{-1}[ISPA]^{-1}[IDI]D_{4,r}^{-1}D_5 > 1 \quad \text{(S45)}$$

$$K_{I,FPP}^{-1}[FPP] > 1 \quad \text{(S46)}$$

$$K_{I,FPP}^{-1}K_{M1}[Mev]^{-1}[FPP] > 1 \quad \text{(S47)}$$

$$K_{M2}[MevP]^{-1} > 1 \quad \text{(S48)}$$

$$K_{M3}[MevPP]^{-1} > 1 \quad \text{(S49)}$$

$$K_{M4}[IPP]^{-1} > 1 \quad \text{(S50)}$$

$$K_{M4}[DMAPP]^{-1} > 1 \quad \text{(S51)}$$

$$K_{M5,IPP}^{-1}K_{M5,DMAPP}[IPP][DMAPP]^{-1} > 1 \quad \text{(S52)}$$

$$K_{M5,DMAPP}[IPP][DMAPP]^{-1} > 1 \quad \text{(S53)}$$

$$K_{I,GPP}[GPP]^{-1} > 1 \quad \text{(S54)}$$

$$K_{I,FPP}K_{M6,IPP}K_{M6,GPP}^{-1}[IPP]^{-1} > 1 \quad \text{(S55)}$$

$$K_{I,GPP}K_{M6,IPP}K_{M6,GPP}^{-1}[IPP]^{-1}[GPP]^{-1} > 1 \quad \text{(S56)}$$

$$K_{M7}^{-1}[FPP] > 1 \quad \text{(S57)}$$

*Steady-State Solution*

$$[MevP]_{ss} = 3k_{cat2}^{-1}k_{cat7}K_{M2}[PMK]^{-1}[ADS] \quad \text{(S58)}$$

$$[MevPP]_{ss} = 3k_{cat3}^{-1}k_{cat7}K_{M3}[PMD]^{-1}[ADS] \quad \text{(S59)}$$

$$[IPP]_{ss} = k_{cat5}^{-1}k_{cat7}K_{M5,DMAPP}[ISPA]^{-1}[ADS] \quad \text{(S60)}$$

$$[DMAPP]_{ss} = k_{cat5}^{-1}k_{cat7}K_{M5,DMAPP}[ISPA]^{-1}[ADS] \quad \text{(S61)}$$

$$[GPP]_{ss} = k_{cat6}^{-1}k_{cat5}K_{M5,DMAPP}^{-1}K_{I,GPP}K_{M6,IPP} \quad \text{(S62)}$$

$$[FPP]_{ss} = \frac{1}{3}k_{cat7}^{-1}k_{cat1}K_{M1}^{-1}K_{I,FPP}[ADS]^{-1}[MK][Mev] \quad \text{(S63)}$$

$$[AMO]_{ss} = k_{out}^{-1}k_{cat7}[ADS] \quad \text{(S64)}$$

$$D_1 = \frac{1}{3}k_{cat7}^{-1}k_{cat1}[ADS]^{-1}[MK][Mev] \quad \text{(S65)}$$

$$D_2 = K_{M2} \quad \text{(S66)}$$

$$D_3 = K_{M3} \quad \text{(S67)}$$

$$D_{4,f} = K_{M4} \quad \text{(S68)}$$

$$D_{4,r} = K_{M4} \quad \text{(S69)}$$

$$D_5 = k_{cat5}^{-1} k_{cat7} K_{M5,DMAPP}^{2} [ISPA]^{-1} [ADS] \quad \text{(S70)}$$

$$D_6 = K_{I,GPP} K_{M6,IPP} \quad \text{(S71)}$$

$$D_7 = \frac{1}{3} k_{cat1} k_{cat7}^{-1} K_{M1}^{-1} K_{I,FPP} [MK][Mev][ADS]^{-1} \quad \text{(S72)}$$

$$r_{out} = k_{cat7} [ADS] \quad \text{(S73)}$$

*<u>Phenotypic Boundaries (Redundant inequalities are not listed)</u>*

$$k_{cat4} k_{cat5}^{-1} K_{M4}^{-1} K_{M5,DMAPP} [IDI][ISPA]^{-1} > 1 \quad \text{(S74)}$$

$$\frac{1}{3} k_{cat1} k_{cat7}^{-1} K_{M1}^{-1} [MK][Mev][ADS]^{-1} > 1 \quad \text{(S75)}$$

$$\frac{1}{3} k_{cat1} k_{cat7}^{-1} [MK][ADS]^{-1} > 1 \quad \text{(S76)}$$

$$\frac{1}{3} k_{cat2} k_{cat7}^{-1} [PMK][ADS]^{-1} > 1 \quad \text{(S77)}$$

$$\frac{1}{3} k_{cat3} k_{cat7}^{-1} [PMD][ADS]^{-1} > 1 \quad \text{(S78)}$$

$$k_{cat5} k_{cat7}^{-1} K_{M4} K_{M5,DMAPP}^{-1} [ISPA][ADS]^{-1} > 1 \quad \text{(S79)}$$

$$K_{M5,IPP}^{-1} K_{M5,DMAPP} > 1 \quad \text{(S80)}$$

$$k_{cat5} k_{cat7}^{-1} [ISPA][ADS]^{-1} > 1 \quad \text{(S81)}$$

$$k_{cat6} k_{cat5}^{-1} K_{M5,DMAPP} K_{M6,GPP}^{-1} > 1 \quad \text{(S82)}$$

$$k_{cat5} k_{cat7}^{-1} K_{M5,DMAPP}^{-1} K_{I,GPP} K_{M6,IPP} K_{M6,GPP}^{-1} [ISPA][ADS]^{-1} > 1 \quad \text{(S83)}$$

$$k_{cat6} k_{cat7}^{-1} [ISPA][ADS]^{-1} > 1 \quad \text{(S84)}$$

$$\frac{1}{3}k_{cat1}k_{cat7}^{-1}K_{M1}^{-1}K_{I,FPP}K_{M7}^{-1}[MK][Mev][ADS]^{-1} > 1 \quad \text{(S85)}$$

**4. Resolving Co-dominant Phenotypes**

Co-dominant phenotypes arise when the steady state influxes or effluxes of a given metabolic pool exhibit the same numerical value. In such cases, dominance cannot be determined. Mathematically, co-dominance leads to inconsistent boundaries of the type 1>1, which renders associated phenotypes invalid. To exemplify this situation, consider Fig. S4A, which represents a simplified metabolic network in which IPP interacts with DMAPP to generate GPP. Eqs. S86 to S88 describe the dynamics of this network. For simplicity, all reactions are described by first order kinetics.

$$\frac{dIPP}{dt} = k_0 - k_1[IPP] - k_2[IPP][DMAPP] \quad \text{(S86)}$$

$$\frac{dDMAPP}{dt} = k_1[IPP] - k_2[IPP][DMAPP] \quad \text{(S87)}$$

$$\frac{dGPP}{dt} = k_2[IPP][DMAPP] - k_3[GPP] \quad \text{(S88)}$$

$k_o$ represents the input flux to the network, while all other $k_i$ are rate constants. A null space analysis of the network's stoichiometric matrix (Fig. S4B) reveals that numerical values for the fluxes $F_1$, $F_2$ and $F_3$ will always be identical for any steady state flux distribution (Fig. S4C). As a result, dominance cannot be determined for fluxes consuming IPP, i.e. $F_1$ and $F_2$. Table S7 contains a mathematical characterization of the two phenotypes exhibited by this network. Note that even though S-system equations for each phenotype are slightly different, their steady state equations for IPP, DMAPP and GPP are identical. As expected, an inconsistent boundary is obtained in each case (i.e., 1>1) after introducing the steady state into the dominance conditions. A four-step strategy is employed to handle co-dominant phenotypes:

1. Relax inconsistent boundary conditions arising from co-dominances (i.e., 1>1) by multiplying the left-hand side of all conditions linked to co-dominances by a factor of 2.

This factor can be rationalized by assuming that one of two identical fluxes is larger than the other. As a result, the selected dominant flux will now support twice its original flux. For example, the dominance condition $k_1^{-1}k_2DMAPP > 1$ of phenotype 2 in Table S7 turns into $2k_1^{-1}k_2DMAPP > 1$, which leads to the phenotypic boundary $2 > 1$ and renders phenotype 2 valid.

2. Track the number of co-dominant fluxes affecting each metabolic pool. For phenotype 2, we have two co-dominant effluxes out of the metabolic pool IPP, i.e., $k_1[IPP]$ and $k_2[IPP][DMAPP]$.
3. Adjust S-system equations and corresponding dominance conditions to reflect the existence of independent and/or linked co-dominant fluxes in the network. The adjustment consists in multiplying the *input flux* of the affected metabolic pools by $n$ or $1/n$, with $n$ the number of co-dominant fluxes. The former adjustment is performed if the co-dominant fluxes are influxes and the latter if the co-dominant fluxes are effluxes.
4. Co-dominant fluxes generate overlapping, redundant phenotypes with identical phenotypic boundaries and steady state behavior (Table S7). To facilitate the analysis of biochemical networks containing co-dominant fluxes, we remove redundant phenotypes from the system's phenotypic repertoire.

Table S8 illustrates the application of the previous steps to process phenotype 2. Computational routines in the Design Space Toolbox v.2.0 (Lomnitz and Savageau, 2016) already implemented the first step of this strategy. Here, we expanded existing routines to incorporate steps 2 to 4 to facilitate the analysis of co-dominant phenotypes.

### 5. Effect of Feedback Inhibition on the Global Landscape of Metabolic Engineering Strategies

Assuming that the input flux is not affected by removing the MK feedback inhibition, equivalent operating points for the modified network can be estimated. We first use Equations S89 and S90 to calculate the steady state concentration of FPP and the input flux of the original network, respectively.

$$[FPP]_{ss} = \frac{1}{3} k_{cat}^{-1} k_{cat1} K_{M1}^{-1} K_{I,FPP} [ADS]^{-1} [MK][Mev] = 69.9 \mu M \quad \text{(S89)}$$

$$F_{inSS} = \frac{k_{cat1}[MK][Mev]}{K_{M1} + K_{M1}[FPP]_{SS} K_{I,FPP}^{-1} + [Mev]} = 152.5 \mu M \min^{-1} \quad \text{(S90)}$$

Kinetic parameters for strain mbis3 were used to evaluate both equations. Then, values for $k^*_{cat1}$ and $K^*_{M1}$, which characterize kinetic properties of MK without feedback inhibition, can be calculated using Equation S91:

$$F_{inSS} = \frac{k^*_{cat1}[MK][Mev]}{K^*_{M1} + [Mev]} \quad \text{(S91)}$$

Solving for $k^*_{cat1}$ and evaluating kinetic values for strain mibs3 leads to Eq. S92:

$$k^*_{cat1} = 0.46 \min^{-1} \mu M^{-1} K^*_{M1} + 2.3 \min^{-1} \quad \text{(S92)}$$

Figs. S5C and S5D show the location of two different operating points in Design Space, which are calculated assuming $K^*_{M1}$ values of 131 and 1.31 μM, respectively. Corresponding $k^*_{cat1}$ values are 62.56 and 2.90 $min^{-1}$. The phenotype containing the operating point is located within region II and III, respectively.

### 6. Supplementary Tables

**Table S1. Experimentally Determined Protein Concentrations for Three Strains.** These values were determined by Weaver et. al. (2015) and are used, along with the kinetic parameters listed in Table S2, to define the operating points for the strains mbis3 (wild-type), saMK3 and 10kADS. Protein concentrations are given in μM. Enzyme prefixes stand for sc: *Saccharomyces cerevisiae*; sa: *Staphylococcus aureus*; ec: *Escherichia coli* and aa: *Artemisia annua*.

| **Enzyme** | **Strain** | | |
| --- | --- | --- | --- |
| | **mbis3** | **saMK3** | **10kADS** |
| **scMK** | 66.3 | | 197 |
| **saMK** | | 23.2 | |
| **scPMK** | 2.9 | 4.9 | 14.1 |
| **scPMD** | 2.2 | 2.7 | 2.3 |
| **ecIDI** | 18.4 | 15.3 | 21.6 |
| **ecISPA** | 35 | 36.8 | 24.2 |
| **aaADS** | 39.6 | 25.6 | 678 |

**Table S2. Reference Kinetic Parameters Used in this Study.** The kinetic parameters listed here were compiled by Weaver et. al. (2015) and are used, along with experimentally determined protein concentrations (Table S1), to define the operating points for the strains mbis3, saMK3 and 10kADS. $K_{cat}$ values are in units of $s^{-1}$, $K_M$ and $K_I$ values are in units of μM. Note that Weaver et. al. (2015) experimentally determined a $kcat_7$ value of 0.022 ± 0.008 $s^{-1}$. We used the updated $kcat_7$ value for all analyses reported in this work.

| Enzyme | Kinetic Parameter | Value |
|---|---|---|
| *S. cerevisiae* mevalonate kinase (scMK) | $k_{cat1}$ | 38 |
| | $K_{M1}$ | 131 |
| | $K_{I,\ FPP}$ | 1.9 |
| *S. aureus* mevalonate kinase (saMK) | $k_{cat1}$ | 6 |
| | $K_{M1}$ | 41 |
| | $K_{I,\ FPP}$ | 46 |
| *S. cerevisiae* phosphomevalonate kinase (scPMK) | $k_{cat2}$ | 10 |
| | $K_{M2}$ | 885 |
| *S. cerevisiae* mevalonate diphosphate decarboxylase (scPMD) | $k_{cat3}$ | 4.9 |
| | $K_{M3}$ | 123 |
| *S. cerevisiae* mevalonate diphosphate Isomerase (ecIDI) | $k_{cat4}$ | 0.33 |
| | $K_{M4}$ | 8 |
| *E. coli* farnesyl pyrophosphate synthase (ecISPA, reaction 1) | kcat5 | 0.21 |
| | $K_{M5,IPP}$ | 1.3 |
| | $KM_{5,DMAPP}$ | 29.3 |
| *E. coli* farnesyl pyrophosphate synthase (ecISPA, reaction 2) | $k_{cat6}$ | 0.47 |
| | $K_{M6,\ IPP}$ | 10.3 |
| | $K_{M6,\ GPP}$ | 5.5 |
| | $K_{I,\ GPP}$ | 74 |
| *A. annua* Amorphadiene Synthase (aaADS) | $k_{cat7}$ | $6.8x10^{-3}$ |
| | $K_{M7}$ | 3.3 |
| Amorphadiene Export | $k_{out}$ | 1/60 |

**Table S3. Eigenvalue Analysis for Phenotypes 7306_3 and 7330_3.** Eigenvalues were calculated for each biochemical phenotype using NumPy's linear algebra package. The operating point of the network within phenotype **7306_3** (black circle in Fig. 2A) is defined by the experimentally measured protein concentrations for the base strain mbis3 and kinetic parameters extracted from the literature (Tables S1 and S2): MK = 66.3 μM, PMK = 2.9 μM, PMD = 2.2 μM, IDI = 18.4 μM, ISPA = 35 μM, ADS = 39.6 μM, Mev =5 μM, $k_{cat1}$ = 38 $s^{-1}$, $K_{M1}$ = 131 μM, $K_{I,\,FPP}$ = 1.9 μM, $k_{cat2}$ = 10 $s^{-1}$, $K_{M2}$ = 885 μM, $k_{cat3}$ = 4.9 $s^{-1}$, $K_{M3}$ = 123 μM, $K_{M4}$ = 8 μM, $k_{cat4}$ = 0.33 $s^{-1}$, $k_{cat5}$ = 0.21 $s^{-1}$, $K_{M5,IPP}$ = 1.3 μM, $K_{M5,DMAPP}$ = 29.3 μM, $k_{cat6}$ = 0.47 $s^{-1}$, $K_{M6,IPP}$ = 10.3 μM, $K_{M6,GPP}$ = 5.5 μM, $K_{I,GPP}$ = 74 μM, $k_{cat7}$ = $2.2 \times 10^{-2}$ $s^{-1}$, $K_{M7}$ = 3.3 μM, $k_{out}$ = 1 $min^{-1}$. The operating point located within phenotype **7330_3** (white diamond in Fig. 2A) is identical to the one in phenotype **7306_3**, with the exception of the ADS concentration, which corresponds to 109.65 μM (log(ADS) = 2.04).

| Rank | Eigenvalues 7306_3 | | Eigenvalues 7330_3 | |
|---|---|---|---|---|
| | **Real** | **Imaginary** | **Real** | **Imaginary** |
| **1** | -45.542 | 0.000 | -45.540 | 0.000 |
| **2** | -14.946 | 0.000 | -15.791 | 0.000 |
| **3** | -6.083 | 0.000 | -8.717 | 3.386 |
| **4** | -4.265 | 0.000 | -8.717 | -3.386 |
| **5** | -1.0 | 0.000 | -1.000 | 0.000 |
| **6** | -0.738 | 0.897 | 0.295 | 2.570 |
| **7** | -0.738 | -0.897 | 0.295 | -2.570 |

**Table S4. Physiological Phenotypes of the Amorphadiene Network and their Case Signatures.** We used DST3 to enumerate the phenotypic repertoire of the network defined by Eqs. S9 – S24. A total of 40 phenotypes were sorted into four different groups according to their logarithmic gain patterns (see Fig. 3). Groups II and III are analyzed independently to better understand the effect of MK kinetic parameters on pathway flux. For each phenotype, a case signature indicating dominant terms in each equation is provided. For this specific network, dominance signatures consist of 16 two-digit sets of indices, which specify the positive and the negative term dominating in each equation (Eqs. S9 – S24). Phenotypes resulting from cyclical cases contain a three-digit signature, in which the first digit refers to the equation number from which the positive term stems (Valderrama-Gómez et al. 2020).

| Group | Case Number | Case Signature |
|---|---|---|
| I | 6154 | 11 11 13 12 11 11 11 21 11 11 11 11 21 11 21 11 |
| | 6158 | 11 11 13 12 11 11 11 21 11 11 11 11 21 31 21 11 |
| | 7298_3 | 11 11 21 312 11 11 11 21 11 11 11 11 11 11 21 11 |
| | 7302_3 | 11 11 21 312 11 11 11 21 11 11 11 11 11 31 21 11 |
| | 7306_3 | 11 11 21 312 11 11 11 21 11 11 11 11 21 11 21 11 |
| | 7310_3 | 11 11 21 312 11 11 11 21 11 11 11 11 21 31 21 11 |
| | 7330_3 | 11 11 21 312 11 11 11 21 11 11 11 21 21 11 21 11 |
| | 7334_3 | 11 11 21 312 11 11 11 21 11 11 11 21 21 31 21 11 |
| | 7346_3 | 11 11 21 312 11 11 11 21 11 11 21 11 11 11 21 11 |
| | 7350_3 | 11 11 21 312 11 11 11 21 11 11 21 11 11 31 21 11 |
| II | 7681_3 | 11 11 21 312 11 11 11 31 11 11 11 11 11 11 11 11 |
| | 7685_3 | 11 11 21 312 11 11 11 31 11 11 11 11 11 31 11 11 |
| | 7689_3 | 11 11 21 312 11 11 11 31 11 11 11 11 21 11 11 11 |
| | 7693_3 | 11 11 21 312 11 11 11 31 11 11 11 11 21 31 11 11 |
| | 7713_3 | 11 11 21 312 11 11 11 31 11 11 11 21 21 11 11 11 |
| | 7717_3 | 11 11 21 312 11 11 11 31 11 11 11 21 21 31 11 11 |
| | 7729_3 | 11 11 21 312 11 11 11 31 11 11 21 11 11 11 11 11 |
| | 7733_3 | 11 11 21 312 11 11 11 31 11 11 21 11 11 31 11 11 |
| | 6537 | 11 11 13 12 11 11 11 31 11 11 11 11 21 11 11 11 |
| | 6541 | 11 11 13 12 11 11 11 31 11 11 11 11 21 31 11 11 |
| III | 5769 | 11 11 13 12 11 11 11 11 11 11 11 11 21 11 11 11 |
| | 5773 | 11 11 13 12 11 11 11 11 11 11 11 11 21 31 11 11 |
| | 6913_3 | 11 11 21 312 11 11 11 11 11 11 11 11 11 11 11 11 |
| | 6917_3 | 11 11 21 312 11 11 11 11 11 11 11 11 11 31 11 11 |
| | 6921_3 | 11 11 21 312 11 11 11 11 11 11 11 11 21 11 11 11 |
| | 6925_3 | 11 11 21 312 11 11 11 11 11 11 11 11 21 31 11 11 |
| | 6945_3 | 11 11 21 312 11 11 11 11 11 11 11 21 21 11 11 11 |
| | 6949_3 | 11 11 21 312 11 11 11 11 11 11 11 21 21 31 11 11 |
| | 6961_3 | 11 11 21 312 11 11 11 11 11 11 21 11 11 11 11 11 |
| | 6965_3 | 11 11 21 312 11 11 11 11 11 11 21 11 11 31 11 11 |
| IV | 6153 | 11 11 13 12 11 11 11 21 11 11 11 11 21 11 11 11 |
| | 6157 | 11 11 13 12 11 11 11 21 11 11 11 11 21 31 11 11 |
| | 7297_3 | 11 11 21 312 11 11 11 21 11 11 11 11 11 11 11 11 |
| | 7301_3 | 11 11 21 312 11 11 11 21 11 11 11 11 11 31 11 11 |
| | 7305_3 | 11 11 21 312 11 11 11 21 11 11 11 11 21 11 11 11 |
| | 7309_3 | 11 11 21 312 11 11 11 21 11 11 11 11 21 31 11 11 |
| | 7329_3 | 11 11 21 312 11 11 11 21 11 11 11 21 21 11 11 11 |
| | 7333_3 | 11 11 21 312 11 11 11 21 11 11 11 21 21 31 11 11 |
| | 7345_3 | 11 11 21 312 11 11 11 21 11 11 21 11 11 11 11 11 |
| | 7349_3 | 11 11 21 312 11 11 11 21 11 11 21 11 11 31 11 11 |

**Table S5. Conserved Dominant Processes Characterizing Each Phenotypic Group.** This table summarizes conserved dominance pairs for each phenotypic group in Fig. 3 and was obtained by analyzing case signatures in Table S4. Conserved pairs that are common to all groups were not considered. Each one of the 16 dominance pairs in Table S4 can be associated with its respective equation and metabolite pool or enzymatic process. "Exp" stands for amorphadiene export described by Eq. S24.

| Equation | S9 | S10 | S11 | S12 | S13 | S14 | S15 | S16 | S17 | S18 | S19 | S20 | S21 | S22 | S23 | S24 |
|---|---|---|---|---|---|---|---|---|---|---|---|---|---|---|---|---|
| **Pool/ Enzyme** | **MevP** | **MevPP** | **IPP** | **DMAPP** | **GPP** | **FPP** | **AMO** | **MK** | **PMK** | **PMD** | **$IDI_f$** | **$IDI_r$** | **$ISPA_1$** | **$ISPA_2$** | **ADS** | **Exp** |
| **Dominant Processes: Group I** | | | | | | | | 21 | | | | | | | 21 | |
| **Group II** | | | | | | | | 31 | | | | | | | 11 | |
| **Group III** | | | | | | | | 11 | | | | | | | 11 | |
| **Group IV** | | | | | | | | 21 | | | | | | | 11 | |

**Table S6. Biological Interpretation of Characteristic Dominance Signatures**. Dominance signatures can specify either dominant fluxes for a given metabolite pool or dominant processes determining the saturation pattern of a given enzyme. Note that all conserved dominance signatures in Table S5 determine enzyme saturation patterns. An enzyme is said to be saturated if its substrate concentration is larger than its respective $K_M$ value. The enzyme is *not* saturated if the contrary is true. Increasing the flux through a saturated enzyme usually involves increasing its expression level or $k_{cat}$ value. On the other hand, increasing the flux through an unsaturated enzyme can be achieved by any of the previous strategies, as well as by increasing the concentration of its substrates or decreasing its $K_M$ value.

| Group | Characteristic Dominance | Biological Interpretation |
|---|---|---|
| I | S16: 21 | Enzyme MK: The inhibition term $K_{M1}$ * FPP / $K_{I,FPP}$ is the largest. MK is NOT saturated. |
| | S23: 21 | Enzyme ADS: FPP > $K_{M7}$. ADS is saturated. |
| II | S16: 31 | Enzyme MK: Mev is the largest term. MK is saturated. |
| | S23: 11 | Enzyme ADS: $K_{M7}$ > FPP. ADS is NOT saturated. |
| III | S16: 11 | Enzyme MK: $K_{M1}$ is the largest term. MK is NOT saturated. |
| | S23: 11 | Enzyme ADS: $K_{M7}$ > FPP. ADS is NOT saturated. |
| IV | S16: 21 | Enzyme MK: The inhibition term $K_{M1}$ * FPP/$K_{I,FPP}$ is the largest. MK is NOT saturated. |
| | S23: 11 | Enzyme ADS: $K_{M7}$ > FPP. ADS is NOT saturated. |

**Table S7. Characterization of Co-dominant Cases.** The metabolic network shown in Fig. S4A and mathematically described by Eqs. S86 to S88 exhibits two co-dominant phenotypes, which are characterized here. Shown are their case signature, S-system equations, dominance conditions, steady state, and phenotypic boundaries.

| **Case Signature** | Case 1: 11 11 11 | Case 2: 12 11 11 |
|---|---|---|
| **S-system** | $\frac{dIPP}{dt} = k_0 - k_1[IPP]$<br>$\frac{dDMAPP}{dt} = k_1[IPP] - k_2[IPP][DMAPP]$<br>$\frac{dGPP}{dt} = k_2[IPP][DMAPP] - k_3[GPP]$ | $\frac{dIPP}{dt} = k_0 - k_2[IPP][DMAPP]$<br>$\frac{dDMAPP}{dt} = k_1[IPP] - k_2[IPP][DMAPP]$<br>$\frac{dGPP}{dt} = k_2[IPP][DMAPP] - k_3[GPP]$ |
| **Conditions** | $k_1 k_2^{-1} DMAPP^{-1} > 1$ | $k_1^{-1} k_2 DMAPP > 1$ |
| **Steady State** | $IPP = k_0 k_1^{-1}$<br>$DMAPP = k_1 k_2^{-1}$<br>$GPP = k_0 k_3^{-1}$ | $IPP = k_0 k_1^{-1}$<br>$DMAPP = k_1 k_2^{-1}$<br>$GPP = k_0 k_3^{-1}$ |
| **Boundaries** | $1 > 1$ | $1 > 1$ |

**Table S8. Processing Co-dominant Phenotypes.** Co-dominant phenotype 2 of the metabolic network shown in Fig. S4A is processed according to the 4-step strategy presented in Supplementary Section 4. Changes to S-system equations, steady states, dominance conditions and phenotypic boundaries are shown in red. Note that phenotype 1 of the network (see Table S7) is redundant and therefore ignored.

| Step | Equations |
|---|---|
| Relax inconsistent phenotypic boundary conditions (i.e., 1>1) by multiplying the left-hand side of all dominance conditions linked to co-dominances by a factor of 2. | $2k_1^{-1}k_2 DMAPP > 1$ |
| Track the number of codominant fluxes affecting each metabolic pool. Adjust S-system equations and if required, associated dominance conditions. | $\frac{dIPP}{dt} = \frac{1}{2}k_0 - k_2[IPP][DMAPP]$<br>$\frac{dDMAPP}{dt} = k_1[IPP] - k_2[IPP][DMAPP]$<br>$\frac{dGPP}{dt} = k_2[IPP][DMAPP] - k_3[GPP]$ |
| Recalculate steady states and phenotypic boundaries. | Steady States: $IPP = \frac{1}{2}k_0k_1^{-1}$; $DMAPP = k_1k_2^{-1}$; $GPP = \frac{1}{2}k_0k_3^{-1}$<br>Phenotypic boundaries: $2 > 1$ |

### 7. Supplementary Figures

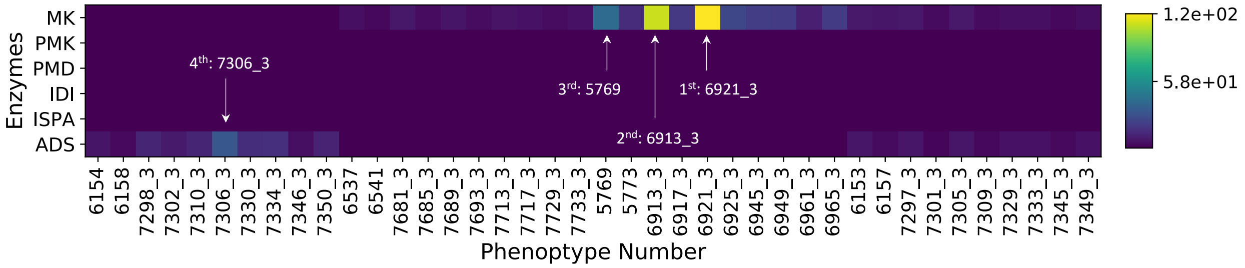


**Figure S1. Phenotypic Landscape Maximizing Production Potential.** Fold change in production flux from a phenotype's nominal operating point is calculated for all valid phenotypes and used as a performance metric to identify top performers. A dark blue color corresponds to a fold change in production flux with a value of 1, meaning that finetuning a given enzyme has no effect on production flux. Numerical values used to generate the heatmap along with a phenotypic ranking can be found in the Supplementary File 2.

| | I | II | III | IV |
|---|---|---|---|---|
| **MK** | **Not Saturated**<br>$K_{M1} * FPP / K_{I,FPP} > Mev$<br>$K_{M1} * FPP / K_{I,FPP} > K_{M1}$ | **Saturated**<br>$K_{M1} < Mev$<br>$K_{M1} * FPP / K_{I,FPP} < Mev$ | **Not Saturated**<br>$Mev < K_{M1}$<br>$K_{M1} * FPP / K_{I,FPP} < K_{M1}$ | **Not Saturated**<br>$K_{M1} * FPP / K_{I,FPP} > Mev$<br>$K_{M1} * FPP / K_{I,FPP} > K_{M1}$ |
| **ADS** | **Saturated**<br>$FPP > K_{M,7}$ | **Not Saturated**<br>$FPP < K_{M,7}$ | **Not Saturated**<br>$FPP < K_{M,7}$ | **Not Saturated**<br>$FPP < K_{M,7}$ |

**Figure S2. Saturation Patterns of Key Enzymes.** An analysis of conserved dominance signatures (Tables S5 and S6) reveals that the saturation pattern of two enzymes, mevalonate kinase (MK) and amorphadiene synthase (ADS), can be used to distinguish between phenotypic groups in Fig. 3.

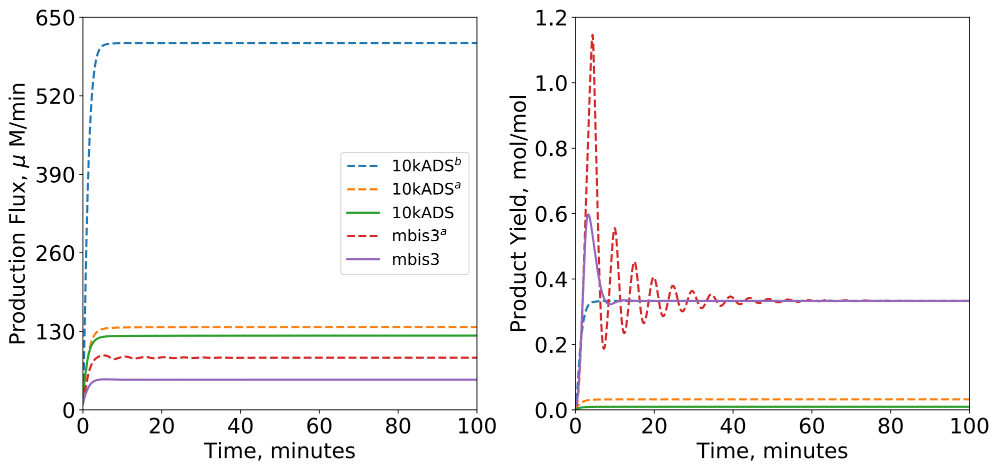


**Figure S3. Production Flux and Yield of Five Different Strains**. Solid lines (green and purple) correspond to model predictions (Eqs. S1 to S8) for experimental strains mbis3 and 10kADS. In each case, the kinetic model was parameterized using values reported in Tables S1 and S2 for the respective strain. Strains mbis3$^a$, 10kADS$^a$, and 10kADS$^b$ are hypothetical strains derived from mbis3. mbis3$^a$ differs from mbis3 solely by its ADS concentration, which corresponds to log(ADS) = 1.8 in mbis3$^a$. Note that this ADS concentration places mbis3$^a$ within phenotype **7306_3** (see Fig. 4A and Fig. 4B). Similarly, strain 10kADS$^a$ differs from mbis3 solely by its ADS concentration, which equals the experimental ADS level of strain 10kADS (678 μM). Finally, strain 10kADS$^b$ is defined by enzyme levels corresponding to the last row of Table 1 in the main text. Note that all strains located within the physiological phenotype **7306_3** (mbis3, mbis3$^a$, and 10kADS$^b$) exhibit a product yield of 1/3, which corresponds to the maximum theoretical value for the network under consideration (Fig. 1). In contrast, the product yield of strains 10kADS and 10kADS$^a$, both located within a pathological region of Design Space, is predicted to have a very low value. This is caused by metabolic imbalances in the network, which, in this specific case, cause the accumulation of MevP, MevPP and IPP (Fig. 2D), and effectively reduce the carbon flux available for production.

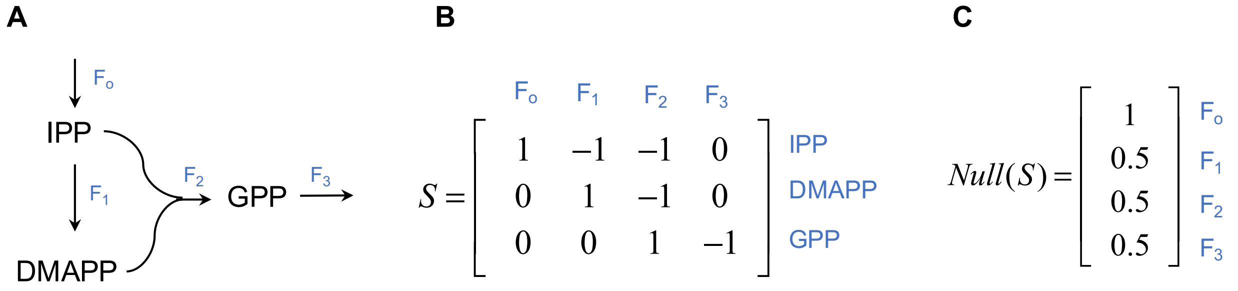


**Figure S4. Simple Network Motif Exhibiting Two Co-dominant Phenotypes. A.** A metabolic network describing a simplified set of reactions between metabolites IPP, DMAPP and GPP is shown. $F_i$ denotes a reaction flux **B.** The network's stoichiometric matrix, S, consists of 3 rows and 4 columns. **C.** The null space of S is shown. The dimension of the null space of S is dim(*Null*(S)) = 1.

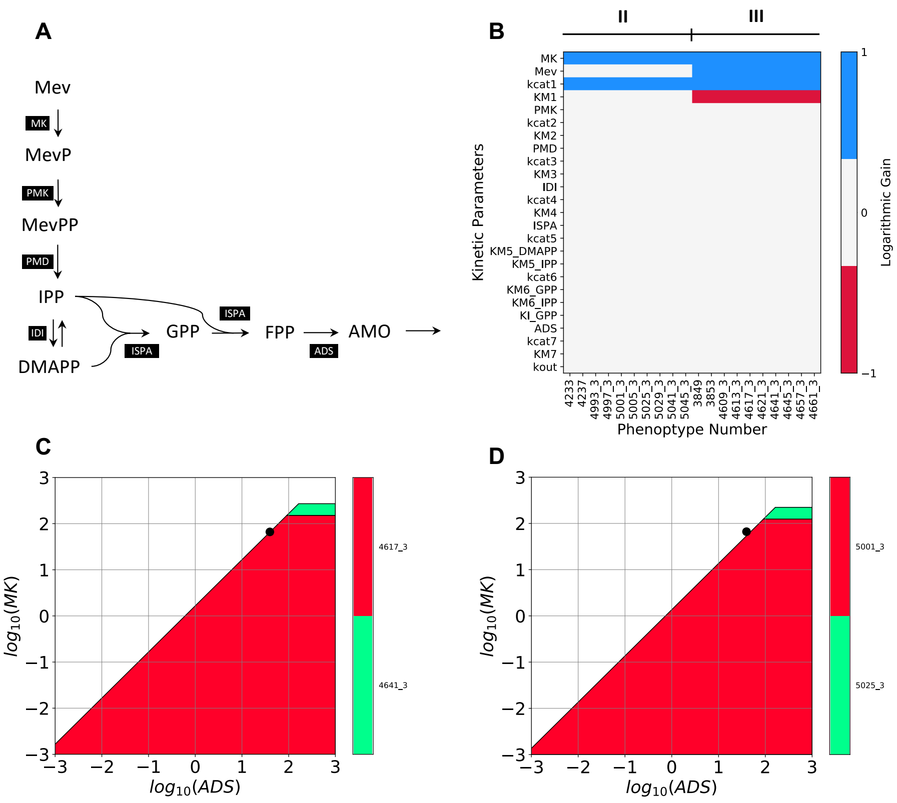


**Figure S5. Effect of Feedback Inhibition on Beneficial Metabolic Engineering Strategies. A.** Modified amorphadiene network lacking feedback inhibition. **B.** Global landscape of metabolic engineering strategies increasing amorphadiene flux for the metabolic network in panel **A**. The modified network exhibits 20 physiological phenotypes. **C.** An operating point of the network (black dot) located on the boundary of phenotype **4617_3** and defined by the mbis3 kinetic parameters except for $K^{*}_{M1}$ = 131 μM and $k^{*}_{cat1}$ = 62.56 min$^{-1}$. **D.** An operating point of the network (black dot) located on the boundary of phenotype **5001_3** and defined by the mbis3 kinetic parameters except for $K^{*}_{M1}$ = 1.31 μM and $k^{*}_{cat1}$ = 2.9 min$^{-1}$.
